# Methylation Clocks Do Not Predict Age or Alzheimer’s Disease Risk Across Genetically Admixed Individuals

**DOI:** 10.1101/2024.10.16.618588

**Authors:** Sebastián Cruz-González, Ogechukwu Okpala, Esther Gu, Lissette Gomez, Makaela Mews, Jeffery M. Vance, Michael L. Cuccaro, Mario R. Cornejo-Olivas, Briseida E. Feliciano-Astacio, Goldie S. Byrd, Jonathan L. Haines, Margaret A. Pericak-Vance, Anthony J. Griswold, William S. Bush, John A. Capra

## Abstract

Epigenetic aging clocks based on DNA methylation patterns across the genome have emerged as a potential biomarker for risk of age-related diseases, like Alzheimer’s disease (AD), and environmental and social stressors. However, methylation clocks have not been comprehensively validated in genetically diverse individuals. Here we evaluate a set of first-, second-, and third-generation methylation clocks in 621 AD patients and matched controls from African American, Hispanic, and White cohorts. The clocks are less accurate at predicting age in genetically admixed cohorts compared to the White cohort, especially for those with substantial African ancestry. This decreased accuracy holds in >2,500 individuals of European and African ancestry from three additional datasets. The clocks also fail to consistently identify age acceleration in admixed AD cases compared to controls. To explore potential causes for the lack of generalization of the clocks, we intersected clock CpGs with methylation, germline genetic variants, and methylation QTL (meQTL) data from global populations. We find differential methylation between African and European ancestry individuals is common for clock CpGs. Genetic variants rarely disrupt clock CpGs between populations, but a substantial fraction of clock CpGs have meQTL with significantly higher frequencies in African genetic ancestries. Our results demonstrate that methylation clocks often fail to predict age and AD risk when applied across populations and suggest avenues for improving their portability by considering differences in genetic and epigenetic patterns across human populations.

## 1 Introduction

Biological aging is the progressive accumulation of cellular damage leading to degeneration and organismal death (Aunan et al., 2016). DNA methylation patterns at CpG sites across the genome correlate strongly with the aging process, an effect that has been quantified using statistical models called “methylation clocks” (Jones, Goodman, and Kobor, 2015). The first generation of methylation clocks were trained to predict chronological age from methylation levels at selected CpGs from across the genome (Hannum et al., 2013; Horvath, 2013; Zhang et al., 2019). A second generation of clocks were trained to use methylation levels to predict mortality risk as proxied by a combination of biomarkers of frailty and physiological decline (Levine et al., 2018; Lu et al., 2019). Finally, a third generation of clocks have been trained to predict the rate of aging based on cohorts with longitudinal data on biomarkers of frailty (Belsky et al., 2022).

Greater predicted DNA methylation age compared to an individual’s chronological age, known as methylation age acceleration, has been associated with an increased risk of many age-related diseases, including coronary heart disease, white matter hyperintensities, Type 2 diabetes mellitus, Parkinson’s disease, and Alzheimer’s disease (AD) (Hodgson et al., 2017; Horvath and Ritz, 2015; Horvath et al., 2016; Levine et al., 2015, 2018; Lu et al., 2019; Raina et al., 2017). As such, methylation clocks show potential as predictive biomarkers of the aging process and age-related health outcomes, and may capture relevant biological signals associated with aging. The clocks are also increasingly being used in social epidemiology research to quantify associations of methylation aging with exposure to adverse social and environmental factors that often differ across groups (Aiello et al., 2024; Chiu et al., 2024; Krieger et al., 2024; Non, 2021).

While methylation is shaped by the environment of an individual, it is also strongly influenced by genetic variation (Kader and Ghai, 2017). Millions of methylation quantitative trait loci (meQTLs)—genetic variants that associate with the methylation level of a CpG site across individuals—have been identified (Smith et al., 2014). MeQTL influence methylation levels via many mechanisms, including disruption of CpGs and effects on transcription factor binding, gene expression, and other gene regulatory processes (Banovich et al., 2014; Oliva et al., 2023). Methylation patterns also vary between human groups, and approximately 75% of variance in methylation between human groups associates with genetic ancestry (Galanter et al., 2017). This suggests that methylation levels and meQTLs often vary in frequency in different genetic ancestries.

Despite these factors that lead to differential methylation levels in different genetic ancestries, the sociodemographic characteristics of the participants whose data were used to construct multiple commonly used methylation clocks is not known (Watkins et al., 2023). Most methylation clock studies do not report population, ancestry, or even geographic descriptors. Given that available genomic data are strongly biased towards individuals of European ancestry (Popejoy and Fullerton, 2016; Sirugo, Williams, and Tishkoff, 2019), we hypothesized that lack of genetic diversity in the training data of methylation clocks could limit their generalizability across global and admixed populations. Similar factors have posed challenges for the application of polygenic risk scores (PRS) across human groups; PRS models often rapidly decrease in accuracy when applied to individuals not represented in the training set (Martin et al., 2019; Novembre et al., 2022; Privé et al., 2022).

To quantify whether current methylation clocks are generalizable across global populations, we analyzed data from MAGENTA, a diverse AD study which has generated blood methylation and genotyping data for 621 individuals from the Americas, including genetically admixed individuals from African American, Puerto Rican, Cuban, and Peruvian cohorts. We evaluated the accuracy of first-, second-, and third-generation methylation clocks at predicting age in these individuals, along with three replication cohorts of African ancestry and European ancestry individuals. We then evaluated whether age acceleration metrics from these clocks associate with AD risk, as they do in individuals of European ancestry. Then, to investigate factors that influence clock performance, we quantified methylation levels, genetic variant frequency, and meQTL patterns for clock CpGs across genetic ancestries. Our results highlight obstacles to the application of methylation clocks as biomarkers for precision medicine and epidemiology, but they also identify promising avenues for considering genetic diversity in the development, application, and interpretation of methylation clocks.

## 2 Results

Our primary analyses are based on genotyping and blood DNA methylation data from 621 individuals from the Americas with AD and non-demented controls collected by the MAGENTA study and >2,500 individuals from three replication cohorts. The MAGENTA study is focused on late-onset Alzheimer’s and thus the average age of participants is 76 years old. Reflecting AD prevalence, the study is 68% female. The individuals come from five cohorts, collected from the United States (White, African American, Cuban), Peru, and Puerto Rico (**Table 1**). We performed replication analyses on whole blood methylation data from African American individuals from the Grady Trauma Project (n = 422) and the GENOA study (n = 1,394), and White Swedish individuals (n = 729) that participated in the Northern Sweden Population Health Study (NSPHS). As described in detail in the Methods, to facilitate comparisons relevant to understand global differences in our study, we use a combination of geographic and race-based identifiers that are likely to best reflect underlying differences in genetic ancestry and admixture components.

**Table 1:** Demographics of the MAGENTA study cohorts.

|  | Alzheimer’s (N=313) | Control (N=308) | Overall (N=621) |
| --- | --- | --- | --- |
| **COHORT** |  |  |  |
| African American | 98 (31.3%) | 107 (34.7%) | 205 (33.0%) |
| Cuban | 22 (7.0%) | 21 (6.8%) | 43 (6.9%) |
| White | 68 (21.7%) | 65 (21.1%) | 133 (21.4%) |
| Peruvian | 41 (13.1%) | 41 (13.3%) | 82 (13.2%) |
| Puerto Rican | 84 (26.8%) | 74 (24.0%) | 158 (25.4%) |
| **SEX** |  |  |  |
| Female | 206 (65.8%) | 213 (69.2%) | 419 (67.5%) |
| Male | 107 (34.2%) | 95 (30.8%) | 202 (32.5%) |

We apply a range of first-, second-, and third-generation methylation-based methylation clocks to these individuals. We then evaluate their accuracy in predicting chronological age, quantify whether they identify accelerated aging in individuals with AD, and explore genetic factors that may influence clock performance (**Figure 1**).

**Figure 1:**
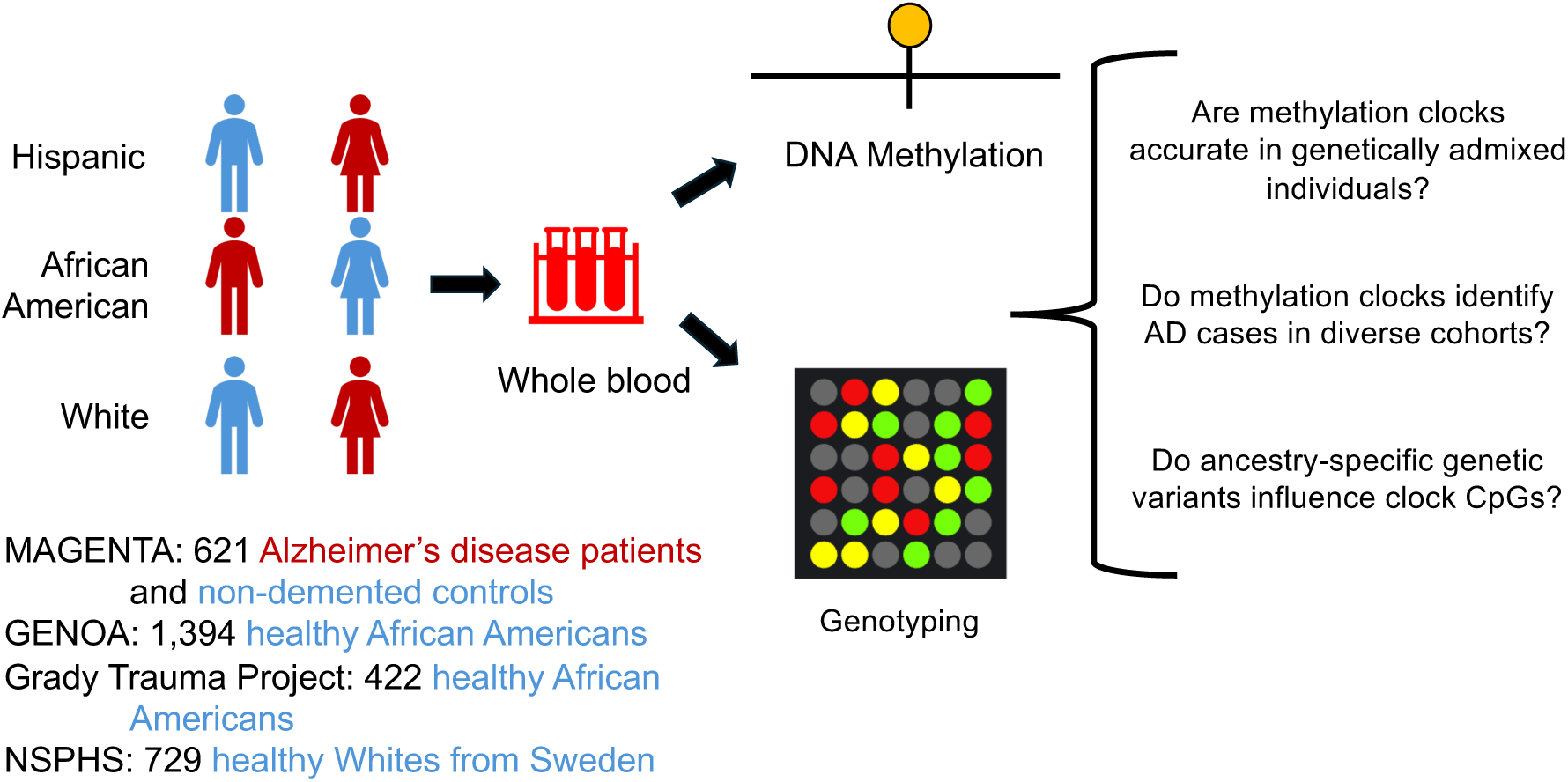
Schematic of the workflow of the study. We analyzed genome-wide methylation and genotyping data from blood samples from 621 AD and non-demented control individuals from the MAGENTA study. To replicate our findings, we analyzed data from 1,394 healthy African Americans from the GENOA study, 422 healthy African Americans from the Grady Trauma Project, and 729 healthy Whites from the NSPHS study. We applied a set of first-, second-, and third-generation methylation clocks to the individuals and estimated their genetic ancestry. This enabled us to explore the performance of methylation clocks in individuals with different genetic ancestries.

### 2.1 Methylation clock accuracy is lower in cohorts with substantial African ancestry

To test whether current methylation clocks are able to predict age accurately in diverse, genetically admixed groups, we evaluated age predictions for the control individuals in the MAGENTA study. We first analyzed the widely used Horvath clock, which was trained on data from several tissues and cell types to accurately predict age across the lifespan using methylation levels at 353 CpG sites.

Age predicted from DNA methylation (DNAm age) in the White cohort using the Horvath clock was strongly correlated with chronological age (Pearson *r* = 0.72) (**Figure 2A**). While this correlation is lower than reported in the original study (>0.9), it is consistent with previous studies of older individuals (Horvath, 2013; Marioni et al., 2015).

**Figure 2:**
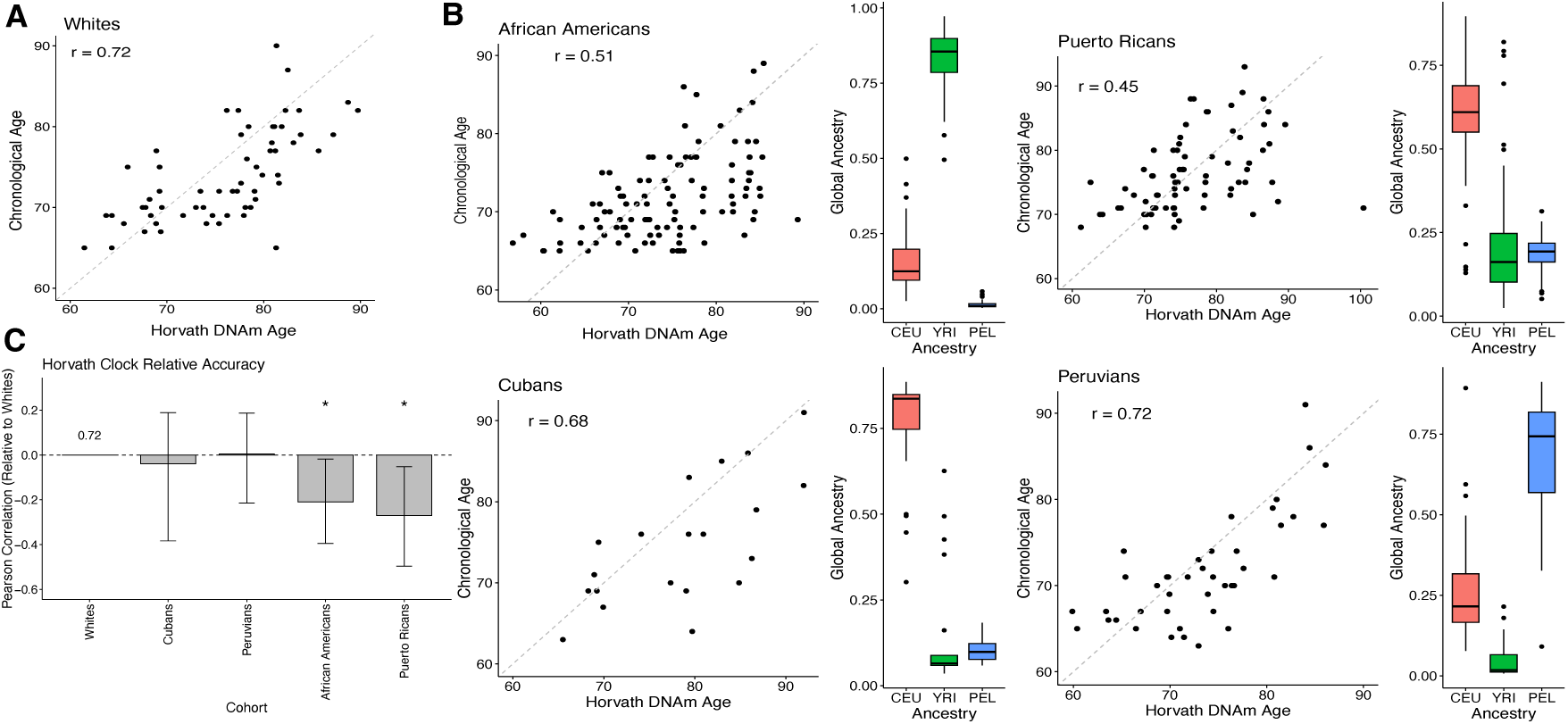
Methylation clock accuracy is lower in cohorts with substantial African genetic ancestry. **A**: Pearson correlation between chronological age and DNAm age predicted by the Horvath clock for controls in the White MAGENTA cohort. The correlation of *r* = 0.72 is consistent with previous studies of similar age groups. **B**: Pearson correlation between chronological age and DNAm age predicted by the Horvath clock for the genetically admixed cohorts in MAGENTA. For each cohort, the adjacent boxplots display the distribution of global ancestry proportions: European (CEU, red), African (YRI, green), and Amerindigenous (PEL, blue). The cohorts with substantial African ancestry—African Americans and Puerto Ricans—exhibit lower correlations (*r* = 0.51 and 0.45, respectively) compared to the Cubans (*r* = 0.68) and Peruvians (*r* = 0.72). **C**: Relative accuracy of the Horvath clock across cohorts compared to Whites. The bar plot shows the difference in Pearson correlation coefficients relative to the White cohort baseline (*r* = 0.72). Asterisks indicate a statistically significant difference from the baseline (* *p <* 0.05).

In comparison to the White cohort, the correlation between DNAm age and chronological age was significantly lower for Puerto Ricans (*r* = 0.45, *p* = 0.007, *n* = 74) and African Americans (*r* = 0.51, *p* = 0.016, *n* = 107) (**Figure 2B**). The correlations for Cubans (*r* = 0.68, *p* = 0.385, *n* = 21) and Peruvians (*r* = 0.72, *p* = 0.52, *n* = 41) were similar to the White cohort. These differences in clock accuracy were not driven by systematic differences in the age distributions for the cohorts (**Supplementary Figure 1**). The methylation clocks were also more accurate in the White cohort than the African American and Puerto Rican cohorts in terms of median absolute error (MAE) (**Supplementary Table 1**). Combining cases and controls did not qualitatively change the performance relationships; the accuracy remained lower for African Americans and Puerto Ricans relative to the Whites (**Supplementary Figure 3**).

The two cohorts with lower correlations come from regions where individuals often have substantial amounts of African ancestry. To explore if admixture levels associated with the accuracy of the Horvath clock in predicting age, we estimated the global proportions of African (YRI), European (CEU), and American (PEL) ancestries in each individual from the MAGENTA cohort using reference groups from the 1000 Genomes Project (The 1000 Genomes Project Consortium et al., 2015).

Methylation clock accuracy was lowest for the cohorts with substantial African ancestry: African Americans (median 85% African) and Puerto Ricans (median 15% African). In contrast, the clocks performed similarly to the White cohort in groups with the lowest African ancestry: Cubans (6% African) and Peruvians (2% African) (**Figure 2C**). We regressed the Horvath clock error on the proportions of African ancestry in the genomes of the MAGENTA individuals, adjusting for chronological age. The proportion of African ancestry is significantly associated with increased Horvath clock error (p = 0.039), with an estimate of 1.46 years more error for 100% African ancestry compared to no African ancestry.

### 2.2 Lower methylation clock accuracy in African ancestry individuals replicates in multiple cohorts

To evaluate if the significantly lower accuracy of the Horvath clock age predictions on individuals with substantial African ancestry holds outside of MAGENTA, we analyzed three independent whole blood methylation datasets. Two focused on African American individuals—the Grady Trauma Project (n = 422) (Katrinli et al., 2024) and the GENOA study (n = 1,394) (Shang et al., 2023)—and one focused on White Swedish individuals (n = 729) (Johansson, Enroth, and Gyllensten, 2013). Each study sampled a wide range of chronological ages, including many older individuals. To enable direct comparison with MAGENTA, we first focused on older individuals (≥ 55 years old). As observed in MAGENTA, the Horvath clock had lower accuracy for the African American cohorts (**Figure 3; bottom row**; Grady: *r* = 0.57, GENOA: *r* = 0.67) and higher accuracy for the White Swedish cohort (*r* = 0.80). We then expanded the analysis to all individuals in each cohort. As expected, the overall accuracy of the age predictions improved, but the disparity in performance between cohorts remained (**Figure 3; top row**; Grady: *r* = 0.88, GENOA: *r* = 0.85, Swedish: *r* = 0.97).

**Figure 3:**
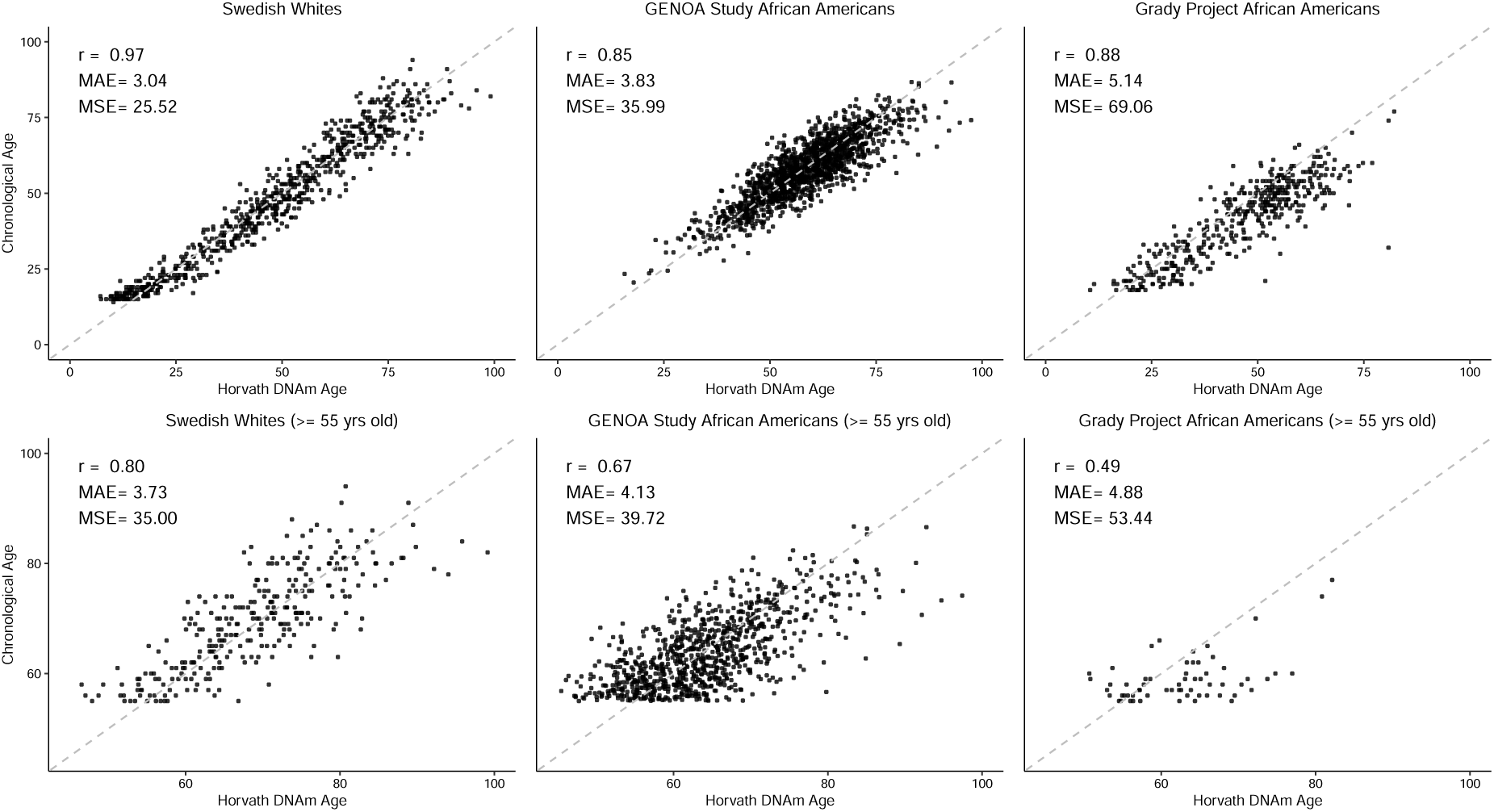
Lower methylation clock accuracy in African Americans holds across cohorts. Scatter plots of chronological age versus Horvath DNAm age in three independent replication cohorts: Swedish Whites (NSPHS), GENOA Study African Americans, and Grady Trauma Project African Americans. **Top Row**: Age predictions for the full age range of each cohort. The Swedish White cohort exhibits the highest accuracy (*r* = 0.97, MAE = 3.04), while both African American cohorts show lower correlations (*r* = 0.85 and *r* = 0.88) and higher error rates (MAE = 3.83 and 5.14, respectively). **Bottom Row**: Age predictions restricted to individuals ≥ 55 years old, similar to the demographics of the MAGENTA cohorts. In this age-restricted subset, the disparity in clock performance is consistent, with the Swedish White cohort maintaining a significantly higher correlation (*r* = 0.80) compared to the GENOA (*r* = 0.67) and Grady (*r* = 0.49) African American cohorts.

A full summary of the clock accuracy statistics, including MAE, maximum absolute error, correlation, and sample size for each cohort are included in **Supplementary Table 2**. Taken together, these results show that the reduced clock accuracy for individuals with substantial African ancestry is not limited to the cohorts in the MAGENTA study, but rather is a general pattern in the portability of methylation clocks.

### 2.3 Accuracy of age prediction on admixed individuals varies across methylation clocks

To investigate the performance of other methylation clocks at predicting chronological age in admixed individuals, we selected several additional publicly available open-source clocks. We considered two other “first-generation” clocks that were trained to predict chronological age: the Hannum clock (Hannum et al., 2013) and a model developed by Zhang et al., 2019 that used large datasets for training and achieved substantially higher performance than previous age predictors. We hereafter refer to this elastic net model as “Zhang_EN”.

Both models achieved higher correlations with chronological age than the Horvath clock across the cohorts in the MAGENTA study. For example, Hannum has a correlation of 0.74 in the White cohort, and consistent with previous evaluations, Zhang_EN has a correlation of 0.88. These relative performance trends held across cohorts, but again, the cohorts with substantial African ancestry, African Americans and Puerto Ricans, consistently had the lowest age correlations for each clock (**Figure 4**).

**Figure 4:**
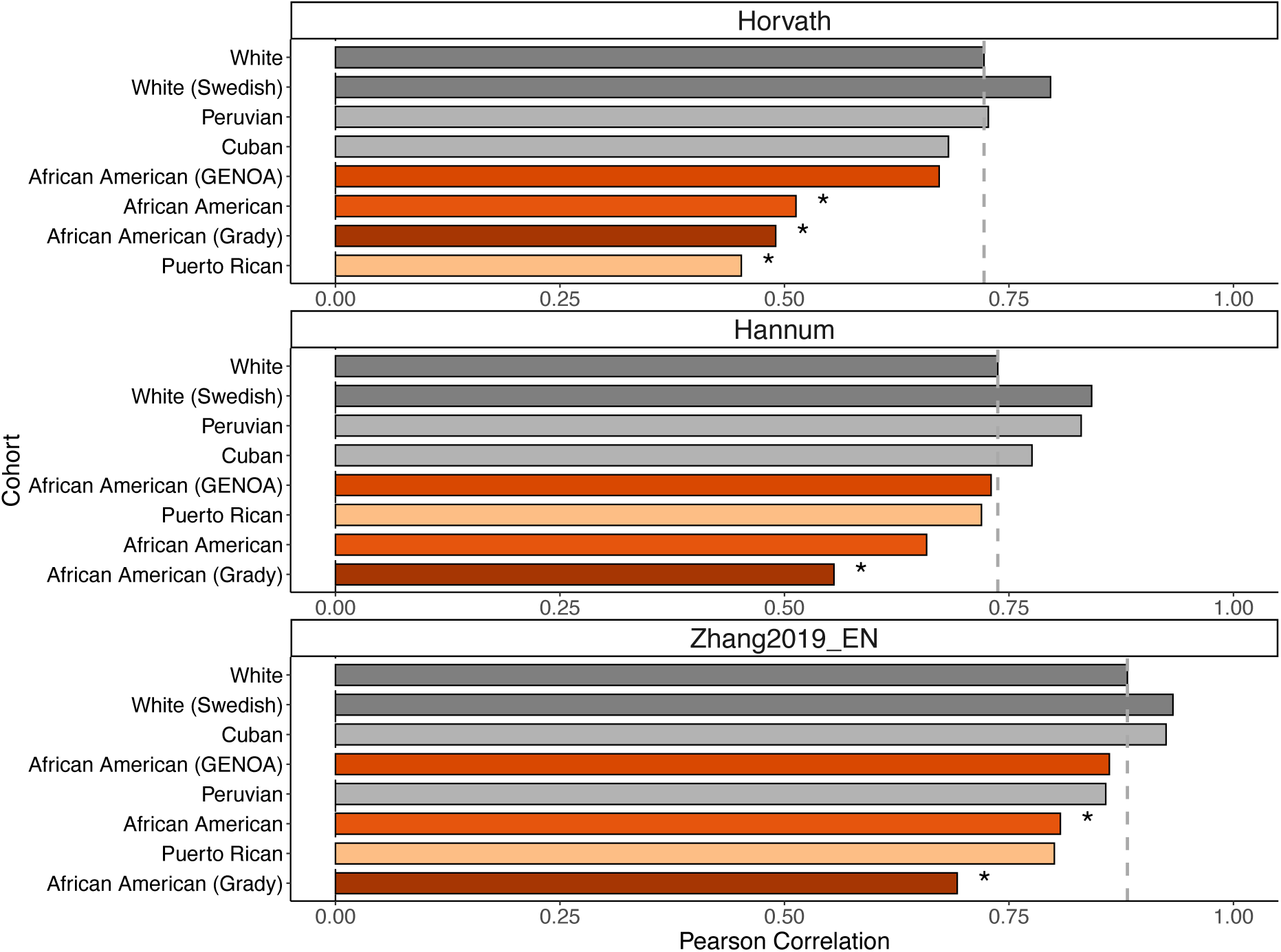
Lower methylation clock accuracy in cohorts with African ancestry holds across multiple clocks. Pearson correlation coefficients between chronological age and DNAm age predicted by the **Horvath** (top), **Hannum** (middle), and **Zhang_EN** (bottom) clocks across all MAGENTA and replication (GENOA, Grady, Swedish) cohorts. The vertical dashed line in each panel represents the baseline correlation observed in the MAGENTA White cohort. Cohorts with substantial African ancestry (African American and Puerto Rican, red bars) consistently exhibit lower correlations compared to Whites, Peruvians, and Cubans across all three clock models. Asterisks indicate a statistically significant difference in correlation compared to the White MAGENTA cohort baseline (* *p <* 0.05).

Next, we evaluated the PhenoAge clock, a “second-generation” clock that is trained on biomarkers of frailty and physiological deterioration (Levine et al., 2018). The correlation between DNAm age and chronological age was lower for this clock in comparison to the other methylation clocks (*r* = 0.53 in the White cohort). This is likely due to the fact that this clock was not trained to predict age directly, but rather markers of aging. This clock did not show as substantial a difference in performance between cohorts as seen for the Horvath clock, but the African American and Puerto Rican individuals again had the lowest correlation of all cohorts. Overall, these results demonstrate that current methylation clocks vary in the correlation of their predicted DNAm age with chronological age in genetically admixed cohorts. The clocks are also consistently the least accurate in predicting age in cohorts with substantial proportions of African ancestry.

### 2.4 Principal component versions of the methylation clocks also have lower age prediction accuracy for genetically admixed individuals

Principal component (PC) versions of many methylation clocks have been developed with the aims of reducing noise and improving replicability and generalizability of age predictions (Higgins-Chen et al., 2022). Thus, we evalauted the accuracy and stability of PC methylation clocks across genetic ancestries. Overall, the PC version of the Horvath clock did not result in consistent improvement in age prediction accuracy or generalization across MAGENTA cohorts (**Supplementary Figures 4 and 5**). The lower accuracy for age prediction in individuals of substantial African ancestry were present for the PC versions of the clocks in the replication cohorts, just as in the MAGENTA cohorts (**Supplementary Figure 6**). Taken together, these results indicate that PC-based versions of the existing methylation clocks do not provide generalizable age predictors for individuals of diverse genetic ancestries.

**Figure 5:**
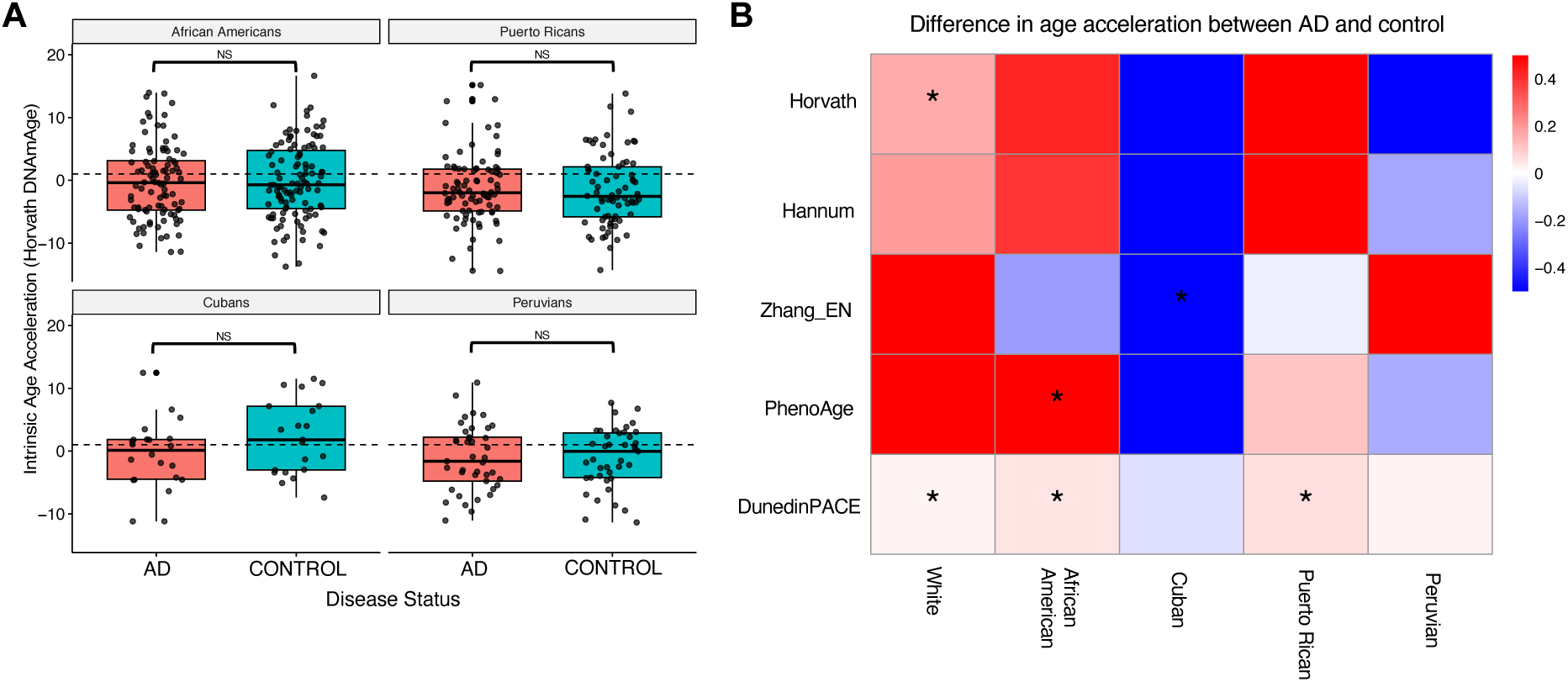
Methylation clocks do not consistently identify accelerated aging in admixed Alzheimer’s cohorts. **A**: Comparison of the distributions of Horvath intrinsic age acceleration for AD patients and non-demented controls for each of the admixed cohorts in MAGENTA. AD patients do not show significantly higher age acceleration in any of the admixed cohorts. In contrast, the AD cases had significantly greater acceleration than controls in the white cohort (**Supplementary** Figure 6). NS = Not significant. **B**: Median differences (in years) in intrinsic age acceleration between AD patients and non-demented controls for five methylation clocks for each cohort in MAGENTA. The clocks do not consistently identfy accelerated aging in AD across cohorts, and the results also vary within cohorts. * *p <* 0.05.

### 2.5 Most methylation clocks do not identify accelerated aging in admixed Alzheimer’s cohorts

DNAm age has been proposed as a promising biomarker and predictive tool for age-related disease risk, particularly because of associations between accelerated DNAm age (compared to chronological age) and the presence of diseases such as coronary heart disease, Parkinson’s disease, and AD. However, these results have largely been observed in European-ancestry cohorts. To evaluate the ability of methylation clocks to identify accelerated aging and risk for age-related disease in diverse, genetically admixed individuals, we quantified the association of methylation age acceleration with AD status in cohorts from the MAGENTA study. In addition to the clocks tested in the previous section, we also included a “third generation” clock, DunedinPACE, that aims to predict the pace of aging as measured by change in biomarkers over time from methylation data, rather than age itself (Belsky et al., 2022).

The cell type composition in blood is known to change with age, which if not accounted for, can confound age acceleration estimates (Jaiswal and Ebert, 2019). Thus, we focused on intrinsic age acceleration estimates computed using established algorithms to correct for cell type composition.

To establish a baseline for this analysis, we tested whether individuals with AD in the White cohort show significantly greater age acceleration than non-demented controls. As expected from previous studies (Levine et al., 2015, 2018), AD cases have modest but significantly greater age acceleration as measured by the Horvath clock than controls (**Supplementary Figure 7**; median 1.7 vs. 1.5 years, *p* = 0.041). For each of the other clocks, AD cases had higher median age acceleration than controls (**Figure 5B**), though the differences only reached statistical significance for the DunedinPACE clock (median 1.09 vs. 1.07, *p* = 0.044).

Having established that previously reported age acceleration in AD was detectable in the MAGENTA White cohort, we evaluated whether the clocks found accelerated aging in the admixed AD cohorts. Focusing first on the Horvath clock, we observed inconsistent relationships between age acceleration and AD status. None of the admixed cohorts showed a significant difference, and controls even had higher median age acceleration in Peruvians and Cubans (**Figure 5A**). Across the other clocks, none consistently identified greater age acceleration in AD cases across all populations (**Figure 5B**). Among the first- and second-generation clocks, only PhenoAge demonstrated a significant ability to differentiate AD cases from controls in any of the non-European ancestry groups, specifically in African American individuals (p = 0.008), which were included in its training set. Cubans consistently showed greater age acceleration in controls rather than cases, while none of the other admixed cohorts even had consistent directions of effect across methods. While sample size in these cohorts is relatively small, our power calculations suggest that low statistical power is unlikely to explain the lack of signal (see Discussion).

DunedinPACE stood out in the evaluation, as it identified significantly greater aging in AD cases compared to controls in the White (p = 0.044), African American (p = 0.0019), and Puerto Rican (p = 0.0090) cohorts using its “pace of aging” metric. However, no significant differences were found for Cubans (p = 0.26) or Peruvians (p = 0.81).

### 2.6 Combining predictions across methylation clocks does not improve their performance

Inspired by recent work on the ensembling of PRS to better predict disease risk from genetic variation across populations (Monti et al., 2024), we evaluated whether combining age predictions could lead to greater accuracy in age prediction and AD risk prediction in the admixed cohorts. To investigate this, we averaged the age predictions for each individual in the MAGENTA study across five methylation clocks: Horvath, Hannum, Zhang_EN, Zhang_BLUP, and PhenoAge clocks. (The Zhang_BLUP is a variation of the Zhang_EN clock that does not use strong regularization.)

The ensemble method’s DNAm age prediction is more strongly correlated with chronological age in comparison to the Horvath and PhenoAge clocks, but it did not improve upon the best predictors (Zhang clocks) across populations (**Supplementary Figure 8A**).

We next evaluated whether the ensemble intrinsic age acceleration estimates would associate more strongly with AD disease status relative to the standalone methylation clocks. Following the same evaluation framework as for the individual clocks, we found only one significant difference in age acceleration. Cuban control individuals had significantly *lower* age acceleration than AD cases (**Supplementary Figure 8B**). Thus, a simple ensemble does not lead to stronger performance at either task in admixed cohorts.

### 2.7 Many clock CpGs are differentially methylated between European and African ancestry individuals

Our results so far demonstrate that existing methylation clocks do not perform consistently across genetically admixed individuals. We now shift our attention to investigating possible mechanisms underlying this lack of generalization. We hypothesized the differential methylation levels between genetic ancestries at clock CpGs could contribute to the lower performance of methylation clocks in admixed individuals. To test for this pattern, we analyzed whole blood methylation data from the EWAS Hub for 306 individuals of African (AFR) and 300 of European (EUR) ancestry. We fitted a linear regression model to the whole blood methylation profiles for these two sets of individuals, including covariates for sex and chronological age. We called CpG sites as differentially methylated in AFR vs. EUR individuals controlling the false discovery rate at 5% with the Benjamini-Hochberg correction. We identified 73,819 differentially methylated CpG sites and intersected them with CpGs included in each clock.

All of the clocks considered had a substantial proportion of their CpGs differentially methylated in AFR vs EUR individuals (**Figure 6A**). The PhenoAge and Horvath clocks had the highest fractions with 24.8% (127/513) and 23.8% (84/353) of their CpG sites differentially methylated, respectively. The other clocks also had substantial fractions of their CpGs differentially methylated: Zhang EN clock (12.6%, 65/514), DunedinPACE (9.2%, 16/173), and the Hannum clock (7%, 5/71). This suggests systematic differences in methylation patterns between genetic ancestries at CpG sites considered in methylation clocks are common.

**Figure 6:**
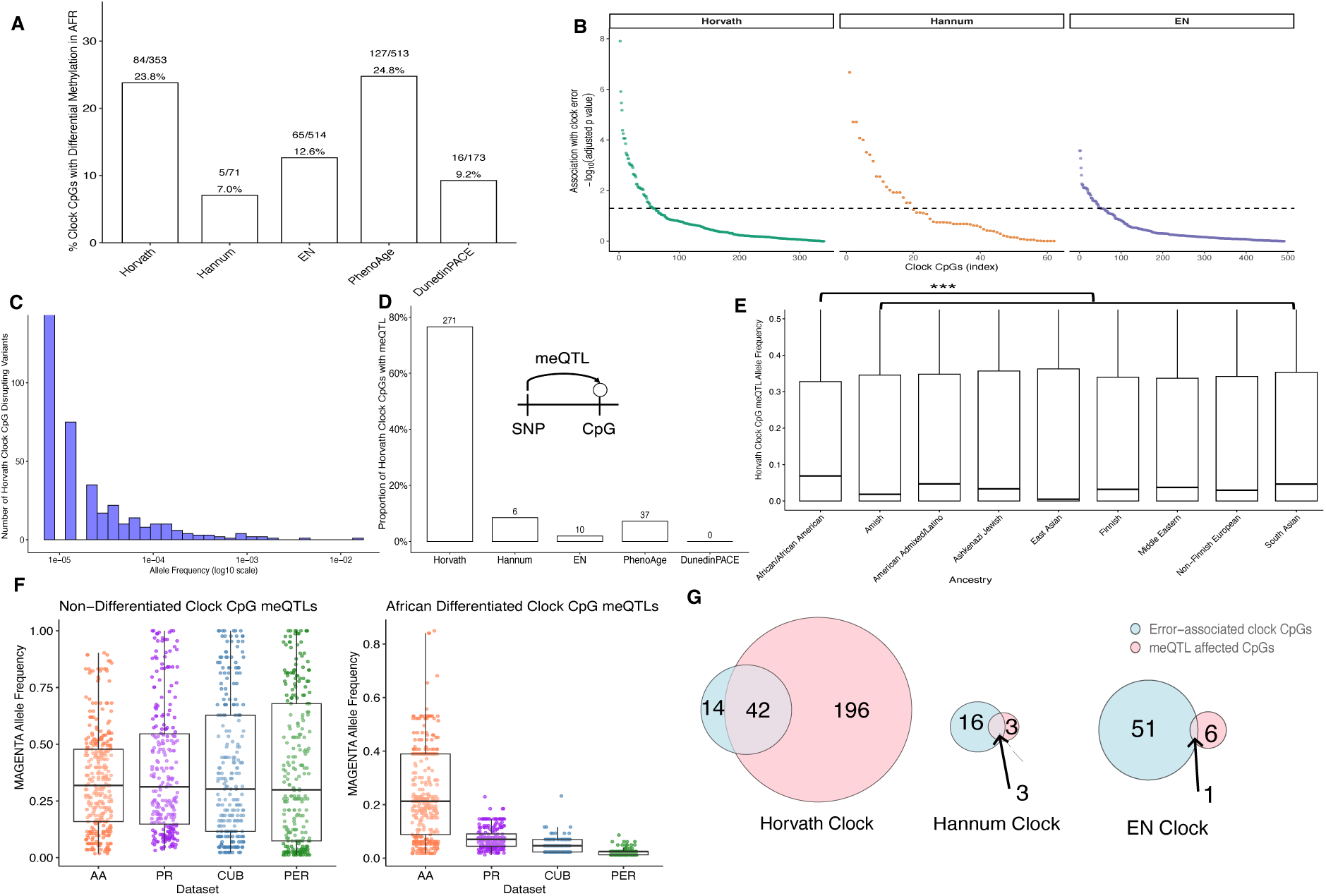
Genetic variants with different frequencies between ancestries associate with methylation levels at clock CpGs. **A**: Percentage of CpG sites in each methylation clock that exhibit differential methylation in whole blood between individuals of African vs. European ancestry. **B**: Methylation levels at some clock CpG sites significantly associate with error across MAGENTA individuals CpGs for each first generation clock are sorted based on the multiple-testing-corected significance of their association with clock error in MAGENTA. **C**: The allele frequency distribution of the single nucletide variants that disrupt CpG sites considered by the Horvath clock in 76,156 individuals from gnomAD (v3.0). Of the 353 clock CpG sites, 245 (69%) have at least one variant, but nearly all the variants are very rare. **D**: Number of CpGs in each clock that are influenced by methylation quantitative trait loci (meQTLs). The Horvath clock contains 271 meQTL-associated CpGs, substantially more than the Hannum, EN, PhenoAge, or DunedinPACE clocks. **E**: Clock meQTL have significantly higher alelle frequency in individuals with African genetic ancestry from gnomAD than all other ancestry groups (median 0.068 for African vs. 0.004–0.046; p < 3.85 × 10*^−^*^25^). **F**: Frequency of meQTL variants in the MAGENTA cohorts. Non-differentiated meQTLs (left) have similar frequencies across cohorts, but African-differentiated meQTLs (right) are significantly more frequent in African Americans (AA) and Puerto Ricans (PR) compared to Peruvians (PER) and Cubans (CUB). **G**: Venn diagrams illustrating the overlap between CpGs that contribute to clock prediction error (“Error-associated”, blue) and those influenced by meQTLs (“meQTL affected”, pink) for the first-generation clocks.

### 2.8 Methylation levels at many CpGs significantly associate with clock error

The presence of differentially methylated CpGs in a clock does not necessarily indicate a barrier to accurate prediction across ancestries. For example, the differential methylation could reflect true differences in environmental exposures between the groups that influence aging. To identify CpGs that could drive the lower accuracy for individuals with substantial African ancestry in their genomes, we tested whether methylation levels at individual clock CpGs are associated with increased clock error across all MAGENTA individuals. The number of CpG sites with methylation levels significantly associated (after Benjamini-Hochberg multiple testing correction) with clock error in the MAGENTA individuals varied across the clocks (**Figure 6B**). The methylation levels at 56 of the 353 Horvath clock CpGs significantly associated with increased clock error, in contrast to 19 of the 71 Hannum clock CpGs. Finally, the methylation levels at 52 of the 514 Zhang EN clock CpGs signficantly associated with increased clock error in MAGENTA individuals. Because the PhenoAge and DunedinPACE clocks are second and third generation clocks, respectively, and therefore do not aim to predict age directly but rather a composite biological age, we did not include these two clocks in this analysis.

### 2.9 Do genetic factors contribute to lower methylation clock performance in admixed populations?

Given that many CpGs sites used in clocks have different methylation levels between genetic ancestries, we sought to investigate potential mechanisms underlying the differenes in methylation and decreased performance of some clocks. Environmental differences between populations likely contribute to the decreased performance of the clocks; however, we do not have comprehensive data on differences in environmental exposures between the cohorts. Moreover, several observations suggest that genetic factors also play a substantial role in clock generalizability. First, genetic ancestry explains a substantial fraction of methylation differences between human populations (Galanter et al., 2017). Furthermore, the particularly large decrease in performance for individuals with a substantial fraction of African ancestry in different populations suggests that genetic divergence rather than the presence of environmental differences alone is responsible.

Thus, we now focus on evaluating potential mechanisms by which genetic variation could reduce clock accuracy across human groups. First, we evaluate whether genetic variation in human populations disupts the potential for methylation at CpG sites. We then explore how often genetic variants that associate with methylation levels (meQTL) influence clock CpGs and differ in frequency between genetic ancestries.

### 2.10 Many methylation clock CpGs are disrupted by genetic variants, but the variants are extremely low frequency

We first quantified how often a genetic variant segregating in human populations disrupted a CpG site included in a clock. This scenario could lead to errors if the variant has different frequencies across cohorts given the loss of potential for methylation and the ability of the site to contribute to the age prediction.

Of the 353 CpG sites considered in the Horvath clock, 245 (69%) have at least one disruptive genetic variant observed in at least one individual in the gnomAD database of variants identified in a cohort of 76,156, including thousands of individuals of non-European ancestry. While this may seem like a substantial concern, these variants are extremely rare both globally and stratifying by ancestry (**Figure 6C; Supplementary Figure 9**). The average frequency is 0.0001, with the most common case being a variant observed in just one individual. Only one clock CpG disrupting variant had a frequency greater than 1%.

The genetic variant frequency patterns are similar for the other clocks (**Supplementary Figure 10**). Of the 71 CpG sites considered in the Hannum clock, 54 (76%) have at least one disruptive variant from the gnomAD database, but only one of the variants has a frequency greater than 1%. For the 514 CpG sites that make up the Zhang19 EN clock, 403 (78%) are disrupted by at least one gnomAD variant, and five of the variants in clock CpGs have a frequency greater than 1%. The PhenoAge clock has 381 of its 513 (74%) disrupted by a gnomAD variant, but only two of these variants have an allele frequency greater than 1%. Finally, in the case of DunedinPACE clock and its 173 CpG sites, 158 (91%) are disrupted by at least one gnomAD variant. However, only one of these variants has an allele frequency greater than 1%. Thus, genetic variation in clock CpG sites themselves is unlikely to be the main cause of the lack of generalization of the methylation clocks.

### 2.11 Common methylation QTL influence clock CpGs

We next assessed the prevalence of meQTLs, genetic variants that associate with clock CpG site methylation, another potential modifier of methylation levels that could lead to spurious DNAm age predictions across individuals. We gathered three sets of meQTLs from Europeans, South Asians, and African Americans (Methods). We intersected the CpGs associated with the meQTLs with clock CpG sites.

Out of the 353 CpGs included in the Horvath clock, 271 (77%) had at least one meQTL. Overall, a total of 29,033 unique variants associated with methylation levels at Horvath clock CpGs. In contrast to CpG disrupting variants, the meQTL had an average allele frequency of 0.19, and 26,500 were common, as determined from gnomAD version 4.1 (global allele frequencies ≥1%). However, only a small proportion of CpG sites in the other first-generation clocks have meQTL (Hannum: 8%, Zhang_EN: 1%). PhenoAge is similar with 7% of its CpGs affected by at least one meQTL. Finally, DunedinPACE had no meQTLs affecting its 173 clock CpG sites (**Figure 6D**).

Thus, the clock with the largest decrease in performance in admixed cohorts (in terms of predicting chronological age and identifying age acceleration in AD) has the largest fraction of CpGs with meQTLs. In contrast, DunedinPACE, the best performing clock at identifying AD cases in the MAGENTA study, had no meQTLs. The three other clocks with intermediate performance in the admixed cohorts, all have meQTL for some CpGs, but much lower fraction than the Horvath clock.

### 2.12 Clock CpG methylation QTLs vary in frequency across ancestries

The presence of meQTL influencing clock CpGs does not necessarily indicate a problem for clock generalization. However, differences in the presence or frequency of meQTL that influence clock CpGs between genetic ancestries could lead to decreased DNAm age prediction accuracy (and therefore weaker associations with disease) in admixed cohorts. For example, if a clock is trained on a cohort without an meQTL, the learned weights for the CpG will not have accounted for the effects of the meQTL. To quantify whether differences in meQTL across genetic ancestries could influence methylation clocks, we analyzed the gnomAD allele frequencies for the 29,033 Horvath clock meQTL tag variants in multiple global populations.

The Horvath clock meQTLs are at significantly higher frequencies in African ancestry populations (median 0.068) than in each of the eight other population groups considered (**Figure 6E**; 0.004–0.046, p < 3.85 *×* 10*^−^*^25^). There were also 2,328 meQTL that were only observed in African individuals.

To connect these results to individuals with recent admixture, like many in the MAGENTA study, we also tested whether the Horvath clock meQTL differed in frequency in local ancestry blocks of different origins in genetically admixed individuals from gnomAD. We used pre-computed local ancestry calls for 7,612 Latino/Admixed American individuals to compare allele frequencies for each meQTL in three ancestral backgrounds: African, Amerindigenous, and European. The clock CpG-affecting meQTLs were at higher frequencies in African local ancestry backgrounds (**Supplementary Figure 12**) relative to Amerindigenous and European backgrounds, consistent with our finding that these meQTLs are most frequent in African ancestry individuals at the global population level.

### 2.13 Differentiated meQTL are present in the MAGENTA cohort

To explore if meQTL influence CpG methylation relevant to clock predictions in the MAGENTA individuals, we tested whether meQTLs described in the previous section are present in MAGENTA. For example, an meQTL (chr3:51639064 A G) for a Horvath clock CpG with a reported beta of 1.34 and an allele frequency of 2% in gnomAD (version 4.1) is present in MAGENTA individuals (study-wide frequency = 11%; AA frequency = 20%; PR frequency = 7.5%; CUB and PER combined frequency = 6%). In individuals with at least one copy of the variant, the methylation level of the corresponding Horvath clock CpG site was higher, as expected from the reported beta for the meQTL. The median methylation level of the clock CpG site for individuals with at least one copy of the variant was significantly higher than for individuals without the variant (median difference in methylation beta = 0.04; *p* = 1.64 × 10*^−^*^7^). Consistent with our hypothesis that this meQTL could contribute to Horvath clock error, individuals with this meQTL had 0.4 years higher median clock error.

Expanding on this example, we quantified whether differences in meQTL frequency across populations observed in other datasets are present in MAGENTA. The overall frequency of meQTL without substantial differentiation in large databases was similar between the MAGENTA admixed cohorts (Figure 6F, left panel; median variant frequencies: AA = 0.32, PR = 0.31, PER = 0.29, CUB = 0.30). However, as expected, the frequency of AFR-differentiated meQTL was higher in the MAGENTA African American (median = 0.21) and Puerto Rican (median = 0.07) cohorts compared to the Peruvian (median = 0.02) and Cuban (median = 0.04) cohorts (**Figure 6F, right panel**). Thus, the MAGENTA individuals carry many meQTL at different frequencies in African ancestry that influence clock CpG methylation.

Finally, we tested whether the error-associated CpGs in the clocks were also affected by meQTL. Strikingly, of the 56 Horvath clock CpGs that were significantly associated with increased clock error in MAGENTA individuals, 42 were also affected by an meQTL, and nine were affected by African ancestry-differentiated meQTL. This pattern contrasts with the Hannum and Zhang EN clocks, where only three and one of the error-associated CpGs were affected by an meQTL, respectively (**Figure 6G and Supplementary Figure 13**).

## 3 Discussion

Methylation clocks are promising biomarkers of aging and social stress, and as tools for mechanistic studies of diseases related to the aging process. Despite their widespread use in these applications, methylation clocks have not been comprehensively evaluated in diverse human groups. These groups are underrepresented in genetic and genomic databases and underserved in biomedicine in terms of access to and quality of healthcare.

In this study, we sought to evaluate the performance of commonly used methylation clocks in genetically admixed individuals from the Americas in the MAGENTA AD study. We found that most clocks did not predict age as accurately in admixed individuals as in White individuals, especially for cohorts with substantial African genetic ancestry. These results replicated in three external, independent cohorts: one of White individuals from Sweden and two of African Americans. We next found that most methylation clocks could not consistently distinguish AD patients and non-demented controls based on age acceleration metrics in non-White cohorts with admixed genetic ancestries.

To evaluate potential genetic factors that could contribute to this decrease in performance, we hypothesized that two types of variants could reduce clock accuracy: 1) variants that disrupt clock CpG sites and prevent methylation and 2) meQTLs that influence clock CpG site methylation. Both scenarios could lead to over or under estimates of age, and therefore spurious associations with age-related disease, if they differ in frequency across genetic ancestries. We discovered that 245 of the 353 CpG sites used by the Horvath clock are disrupted in at least one individual in gnomAD, but these variants are extremely rare and thus unlikely to be a major driver of differences between the cohorts. In contrast, thousands of meQTLs from multiple global populations affected the Horvath clock CpG sites. Many of these meQTL are common, and they are more frequent in individuals with African ancestry in both gnomAD and MAGENTA. We also found that many clock CpGs affected by meQTL also have methylation levels that are significantly associated with clock error in the MAGENTA cohorts. However, we note that the presence of known meQTL was not sufficient to explain all observed clock error at the individual level.

Our findings demonstrate that methylation clocks—a widespread tool in aging, genomics, and social epidemiology research—perform inconsistently across individuals of different genetic ancestries. These results further underline the need for more genetic diversity in the development and evaluation of genomic and epigenomic tools for precision medicine.

We hope that these results also encourage researchers using these tools to exercise caution when interpreting differences in age acceleration. We have shown that many methylation clocks differ significantly in their accuracy at predicting age between cohorts. Thus, what might appear to be a faster pace of aging, could simply be the result of a difference in genetic ancestry from the training cohort. Broadly applying existing methylation clocks could lead to grave consequences and exacerbate existing disparities in access to quality healthcare, as well as provide spurious conclusions about an individual’s health. Due to the increased potential for false positives and false negatives when applying the clocks as predictive biomarkers, individuals at risk might not receive the medical attention they need, and additional stress could unncessarily be placed on individuals in good health. These challenges must be addressed before methylation clocks are adopted as biomarkers for precision medicine.

The challenges we identify here for methylation clocks mirror the limitations of PRS, wherein phenotype prediction models decrease in accuracy on individuals genetically distant from the training population. While there are substantial biological differences in the processes modeled by methylation clocks and PRS, we are optimistic that recent progress on building PRS that are more portable across cohorts will provide strategies for improving methylation clocks.

Our results suggest two promising approaches for building more robust clocks. First, we encourage including individuals from multiple genetic ancestries in the training cohorts. The ability of the PhenoAge clock, which included African Americans in the training cohort, to detect significant age acceleration in the African American AD cases suggests this may improve generalizability. Second, given the large number of meQTL in the human genome and their differences in frequency across human populations, training clocks only on CpG sites that do not have known or population-differentiated meQTL may yield more generalizable clocks. Given that the biological signatures driving methylation clock performance appear to be detectable over large fractions of the genome, removing CpGs with strong meQTL may not substantially limit overall performance. Supporting this, a methylation-based age predictor that is minimally influenced by meQTL performed well in African individuals (Meeks et al., 2025). Moreover, the strong and relatively consistent performance across cohorts of the DunedinPACE clock, which lacks CpGs with meQTL, is also consistent with this approach. It also suggests that methylation clocks that predict the pace of aging (rather than age itself) may be more robust, but further work is needed to validate this hypothesis.

We explored two additional methods for improving generalizability across cohorts that did not yield promising results: ensemble methylation clocks and PC-based methylation clocks. First, motivated by the success of ensembling PRSs for increased accuracy and portability (Truong et al., 2024), we tested whether combining values from multiple methylation clocks could improve risk prediction. For example, combining the predictions of a biomarker-based clock and a first-generation clock could integrate strengths of multiple biomarkers associated with mortality (e.g., PhenoAge) in addition to the chronological aging process and its biological underpinnings (e.g. Horvath clock). However, this approach did not yield improved predictions, though other ensembling approaches could produce different results. We also tested the PC versions of multiple clocks, including the Horvath clock, owing to their reduction in technical noise and variability (Higgins-Chen et al., 2022). However, these clock did not increase accuracy relative to their non-PC versions in the admixed MAGENTA cohorts, and in some cases they performed worse.

There are several caveats and limitations to our study that we hope future work will address. First, the impact of environment on methylation clock accuracy is unresolved. Differences in environmental factors for both local and global populations might lead to decreases in methylation clock accuracy, but they are difficult to study with the data available. Given this, we focused on genetic influences on CpG sites that could lead to spurious associations between populations. The genetic factors we investigated are not sufficient alone to explain differences in clock performance across populations, more work is needed to investigate other non-genetic factors that might cause methylation clocks to not generalize across individuals. Many of the CpGs we discovered with methylation levels associated with clock error do not have known meQTL or variants. We also note that the methods for accounting for cell type composition heterogeneity in blood have not been extensively evalauted across individuals from different populations. Second, while we attempted to evaluate a representative set of first-, second-, and third-generation clocks, we were not able to evaluate all methylation clocks. In particular, some with closed source code that were only available as a web server could not be used due to data privacy restrictions for the MAGENTA samples. Another limitation is the use of blood samples to generate methylation data for the study of a neurodegenerative disease focused on the central nervous system. However, we note that all methylation clocks tested in the present study were developed using blood samples, either exclusively (Hannum, PhenoAge, Zheng EN, Zheng BLUP, and DunedinPACE) or as part of the tissues used in training (Horvath). In addition, these clocks and their association with age-related diseases such as AD have all been validated in multiple tissues, including blood. Multiple studies in the AD literature point to signatures in blood such as gene expression changes, immune cell type composition, and disruption of the blood brain barrier in AD patients relative to non-demented controls, such that signals related to AD pathology can be identified from this peripheral tissue (Griswold et al., 2020; Shigemizu et al., 2022) and through methylation age acceleration (Hodgson et al., 2017; Marioni et al., 2015; Raina et al., 2017).

Finally, while the MAGENTA study is an excellent resource for exploring methylation clocks and AD in admixed individuals, it is not representative of all genetic ancestries and combinations. Moreover, the sample size of the MAGENTA cohort is relatively small. Nonetheless, previous studies have found associations between age acceleration and AD in similar sized cohorts. Both PhenoAge (Levine et al., 2018) and the Horvath clock (Levine et al., 2015) identified age acceleration of less than a year in AD patients relative to non-demented individuals in cohorts of similar sample sizes (e.g., 700 for Levine et al. 2015 and 604 for Levine et al. 2018). We replicated the association with AD in the white MAGENTA cohort, demonstrating the ability to detect methylation effects in MAGENTA. We also performed power calculations based on an effect of the size observed in previous studies (0.5 year acceleration). Stratifying by MAGENTA cohorts, we had 75% power for the African Americans, 72% for the Puerto Ricans, 72% for the Whites, 65% for the Peruvians, and 47% for the Cubans. Thus, it would have been very unlikely to observe no significant differences in any of the admixed cohorts if the effect sizes were similar to previous studies. Thus, reduced clock accuracy is likely a contributor to the lack of association. Furthermore, recent complementary findings on the decreased performance of methylation clocks at age prediction and the role of meQTL in African cohorts (Meeks et al., 2025) further support our conclusions.

In conclusion, our results show that many existing methylation clocks have inconsistent performance and limited portability across genetically admixed cohorts. We encourage future efforts in the development of methylation clocks and other genomics-based aging biomarkers to be genetics- and ancestry-aware to ensure the accuracy of these tools for all individuals, regardless of their genetic background.

## 4 Methods

### 4.1 MAGENTA study

#### Cohort Selection

All participants in the MAGENTA study were recruited through previous studies of AD, including Feliciano-Astacio et al., 2019, Marca-Ysabel et al., 2021, and Griswold et al., 2020. Blood samples were taken for all individuals ascertained and processed at the following sites: the University of Miami Miller School of Medicine (Miami, FL, US), Wake Forest University (Winston-Salem, NC, US), Case Western Reserve University (Cleveland, OH, US), Universidad Central Del Caribe (Bayamón, PR), and the Instituto Nacional de Ciencias Neurologicas (Lima, PE). Ascertainment protocols were consistent across sites and capture cognitive function, family history of AD/related dementias, sociodemographic factors, and dementia staging. All diagnoses were assigned by clinical experts following criteria for diagnosis and staging from the National Institute on Aging Alzheimer’s Association (NIA-AA).

The MAGENTA study is based on pre-existing sample collections which vary in terms of the demographic information collected for each participant. Because the original ascertainment of MAGENTA study participants was international, different population descriptors were used across different ascertainment sites/cohorts. To facilitate comparisons relevant to understand the global differences noted in our study, we use a combination of geographic and race-based identifiers that are likely to best reflect underlying differences in genetic ancestry and admixture components. The label “white” is applied to legacy samples from North Carolina, Tennessee, and South Florida where participants either self-identified with this descriptor or were (in some legacy instances) administratively assigned as White race. The label “African American” is applied to samples collected via ascertainment in North Carolina and South Florida using population descriptors that specifically targeted enrollment of self-identified Black/African American participants. “Puerto Rican”, “Cuban”, and “Peruvian” labels are applied to samples collected as part of ascertainment efforts in these geographic areas. While more precise descriptors of self-identity are preferred, the advanced age of study participants and the older dates of some sample collections make recontact to collect these data impossible. All participants or their consenting proxy provided written informed consent as part of the study protocols approved by the site-specific Institutional Review Boards.

#### Genetic data and ancestry analysis

Genome-wide SNP genotyping was previously performed as previously described for the MAGENTA study cohorts. Briefly, samples were genotyped on the Illumina Infinium Global Screening Array using standard quality control filters on call rate, quality, missingness, and Hardy-Weinberg equilibrium.

For our analyses, local ancestry calls were generated using the *FLARE* software (Browning, Waples, and Browning, 2023) with three reference panels from the 1000 Genomes Project: Utah residents with Northern and Western European ancestry (CEU), Peruvians in Lima (PEL), and Yoruba in Ibadan, Nigeria (YRI). To estimate global ancestry proportions, we summed the haplotype lengths for each ancestry in each individual and divided by the total number of sites considered.

#### Methylation profiles

DNA methylation was quantified using the Illumina HumanMethylation EPICv2.0 according to the manufacturer’s instructions. Samples were randomized across plates and chips to ensure that ancestry, age, and sex were not confounded with each batch. All quality control and data normalization were performed using the the openSeSAMe pipeline from the SeSAMe (Wanding Zhou, 2018) tools for analyzing Illumina Infinium DNA methylation arrays. Probes of poor design were removed from the analysis as well as probes with signal detection P-value >0.05 in more than 5% of the samples. Non-CG probes and probes located on the X, Y, and mitochondrial chromosomes were also removed. Samples with incomplete bisulfite conversion (GCT score >1.5) and principal component analysis outliers were excluded. Noob normalization was performed with SeSAMe, using a nonlinear dye-bias correction. In addition, a principal components analysis of the methylation data was performed. The MAGENTA samples did not stratify by sample plate, cohort, ethnicity, or ascertainment center along any of the first 2 principal components (percentage variance explained for PC1 = 14.96% and PC2 = 6.83%).

### 4.2 Estimating methylation age and its correlation with chronological age in the MAGENTA study

We applied multiple commonly used first-, second-, and third-generation methylation clocks to all individuals in the MAGENTA study with genome-wide methylation data. We used established implementations of the Horvath, Hannum, Zhang_EN, Zhang_BLUP, and PhenoAge clocks from the *methylclock* R library (Pelegí-Sisó et al., 2021). We also applied DunedinPACE, a third-generation clock separately, because it was not included in the *methylclock* library (Belsky et al., 2022). Because this clock does not explicitly predict age, it is not included in the analyses of correlation with biological age. Unless otherwise specified, default options were used for all clocks.

The methylation clocks considered analyze different numbers of CpG sites. For each clock, the sites considered were taken from the *methylclock* library or the original publication. In the case of missing data, the *methylClock* library imputes methylation status using the *mpute.knn* function from the *impute* R library. The MAGENTA cohort had low proportions of missing data for clock CpGs. Specifically, there were 3.7% missing for the Horvath clock, 12.7% for the Hannum clock, 4.5% for the Zhang_EN clock, 3.7% for the PhenoAge clock, and 17.9% for the DunedinPACE clock.

We computed the Pearson correlation of estimated methylation age and chronological age for the controls in each cohort. To compare the strength of correlation between cohorts, we computed p-values for the observed differences using Fisher’s z test and the Zou method for computing confidence intervals as implemented in the *cocor* library (Diedenhofen and Musch, 2015).

### 4.3 Computing methylation age acceleration in Alzheimer’s disease patients and controls

In order to quantify methylation age acceleration from blood methylation data, we estimated raw, intrinsic, and extrinsic age acceleration for all clocks, except DunedinPACE, from the *methylclock* library. Blood cell type composition differs between individuals and over the lifespan; thus, we report results in the main text using intrinsic age acceleration estimates, which capture age acceleration independently of blood cell proportions. We did not have access to empirical estimates of blood cell type counts for the data in MAGENTA, and as such estimated the counts using the functionality for deconvolution implemented in *methylclock*, specifically with the “blood gse35069 complete” parameter. Given that cell type proportions vary between populations based on genetic ancestry, this is an inherent limitation of our study. Nevertheless, the biology of the aging process should be captured using this panel as a reference on our own datasets.

Using the age acceleration estimates, we compared methylation age association between AD cases and matched controls using a Mann-Whitney U test, as implemented in the *stats* library in R. We analyzed the study as a whole and stratified by cohort.

### 4.4 Independent validation cohorts

To evaluate the reproducibility of clock performance across ancestrally diverse populations, we analyzed three independent datasets obtained from the NCBI Gene Expression Omnibus (GEO) using the GEOquery R package. These cohorts were: (1) a European-ancestry cohort from Sweden (GSE87571, *N_total_* = 732), (2) an African American cohort from the Genetic Epidemiology Network of Arteriopathy study (GENOA; GSE210255, *N*_total_ = 1, 394), and (3) an African American cohort from the Grady Trauma Project (GTP; GSE72680, *N*_total_ = 422). Processed methylation beta-value matrices were loaded using the *data.table* package. Probes with high missingness (> 5%) and detection *p*-value columns were removed prior to calculation. To ensure demographic alignment with the MAGENTA study cohort, we also analyzed subsets of each dataset including only individuals aged 55 years or older. This resulted in *n* = 280 for the Swedish cohort, *n* = 865 for the GENOA study, and *n* = 62 for the GTP cohort. DNA methylation age (DNAmAge) was calculated using the *methylclock* package. Clock performance was then quantified using the median absolute error (MAE), mean squared error (MSE), and Pearson correlation between predicted and chronological age.

### 4.5 Evaluating PC methylation clocks

We tested the PC versions of multiple methylation clocks using the *PC Clock* R library (Higgins-Chen et al., 2022; Levine Lab, 2022). We then generated the same accuracy metrics as for the “regular” versions of the clocks when applied to the MAGENTA cohorts and all three replicate cohorts.

### 4.6 Evaluating ensembles of age predictors

We tested the performance of a simple ensemble of age prediction methods at both estimating chronological age and distinguishing AD cases and controls. The ensemble was computed as the average of the estimate of each method on each individual. The resulting predictions were evaluated as described for each clock itself.

### 4.7 Identification of error-associated CpG sites

To identify specific CpG sites whose methylation levels were associated with increased prediction error in the MAGENTA study individuals, we performed a site-wise association analysis across all CpG markers included in the Horvath, Hannum, and Zhang EN clock models. For each clock, we regressed absolute prediction error on the normalized methylation beta values for each individual CpG site within the respective methylation clock. To account for multiple testing, *p*-values were adjusted for each clock separately using the Benjamini-Hochberg (FDR) correction methods. CpG sites were considered significantly associated with prediction error if they had an FDR-adjusted p-value less than 0.05.

### 4.8 Analysis of genetic variation at clock CpG sites

To test whether the CpG sites included in the methylation clocks are variable between individuals, we intersected all the clock CpGs with variants identified in version 3.0 of gnomAD (Karczewski et al., 2020). This database covers 76,156 individuals with whole genome sequencing harmonized from many large-scale sequencing studies. The intersection was performed using *bedtools* in hg38 coordinates (Quinlan and Hall, 2010).

### 4.9 Analysis of meQTL affecting clock CpGs

#### Identification of meQTL affecting clock CpGs

We integrated meQTL sets identified from blood samples by three independent studies. The first study identified 11,165,559 meQTLs from 3,799 Europeans and 3,195 South Asians (Hawe et al., 2022). The second study generated 4,565,687 meQTLs from 961 African Americans (Shang et al., 2023). The final study (EPIGEN) identified 249,710 meQTLs from 2,358 UK individuals (Villicaña et al., 2023). We filtered these sets separately on a false discovery rate threshold of 0.05, correcting for multiple tests using the Benjamini-Hochberg method. These meQTL studies published their results in hg19 coordinates. To integrate with genetic variation and clock CpG data, we mapped the meQTL positions to hg38 using the UCSC liftOver tool (Hinrichs, 2006). For each meQTL set, we intersected the target CpG site with the CpGs considered in each clock and then combined across meQTL sets to generate a set of clock CpGs with evidence of meQTL.

#### Population-level allele frequencies of meQTL affecting clock CpGs

We analyzed the frequency of clock CpG-affecting meQTLs within two different versions of gnomAD. We used version 4.1 to quantify the allele frequencies of these meQTLs in the following global populations: African, Middle Eastern, Admixed American, European (non-Finnish), South Asian, Ashkenazi Jewish, East Asian, European (Finnish), Amish, and a “Remaining” group defined by gnomAD as individuals that did not unambiguously cluster within these previous groups in a principal component analysis. We then used gnomAD version 3.1 to gather allele frequencies for local ancestry blocks identified in 7,612 Latino admixed individuals with varying proportions of European, Amerindigenous, and African ancestry.

## 4.10 Data availability

Raw and normalized beta matrices, along with genotyping data used in this study will be made available at time of publication in the NIAGADS platform. The three validation datasets are publicly available on NCBI GEO. Age predictions for all clocks mentioned in this article will be made available in tab-delimited format in the same Github repository in which all code used for these analyses is available.

## 4.11 Code availability

The publicly available code for analysis are available in the following repository: https://github.com/seba2550/methyl-clocks-admixture

## Acknowledgements

First, we acknowledge the individuals who participated in the MAGENTA study. We are also grateful to members of the Capra Lab, Param Priya Singh, Gillian Meeks, Brenna Henn, and Shyamalika Gopalan for helpful feedback on the project.

This work was supported by the National Institutes of Health (NIH) awards R35GM127087, R01AG070935, U01AG076482, U01AG072579, AG070864, AG072547, and AG074865. It was also supported by a Genentech-UCSF School of Pharmacy Diversity Fellowship to SCG.

This work was conducted in part using the resources of the Wynton High Performance Compute cluster at the University of California, San Francisco.

## 4.12 Author Contributions

Conceptualization: SCG, JAC; Methodology: SCG, JAC; Investigation: SCG, JAC; Resources: EG, LG, MM, JMV, MLC, MRCO, BEFA, GSB, JLH, MAPV, AJG, WSB; Data Curation: SCG, EG, LG, MM, AJG, WSB; Writing – Original Draft: SCG, JAC; Writing - Review and Editing: SCG, AJG, WSB, JAC; Supervision: JAC; Project Administration: AJG, WSB, JAC; Funding Acquisition: SCG, AJG, WSB, JAC.

