## Supplementary Materials for "Methylation Clocks Do Not Predict Age or Alzheimer’s Disease Risk Across Genetically Admixed Individuals"

**Supplementary Figures**

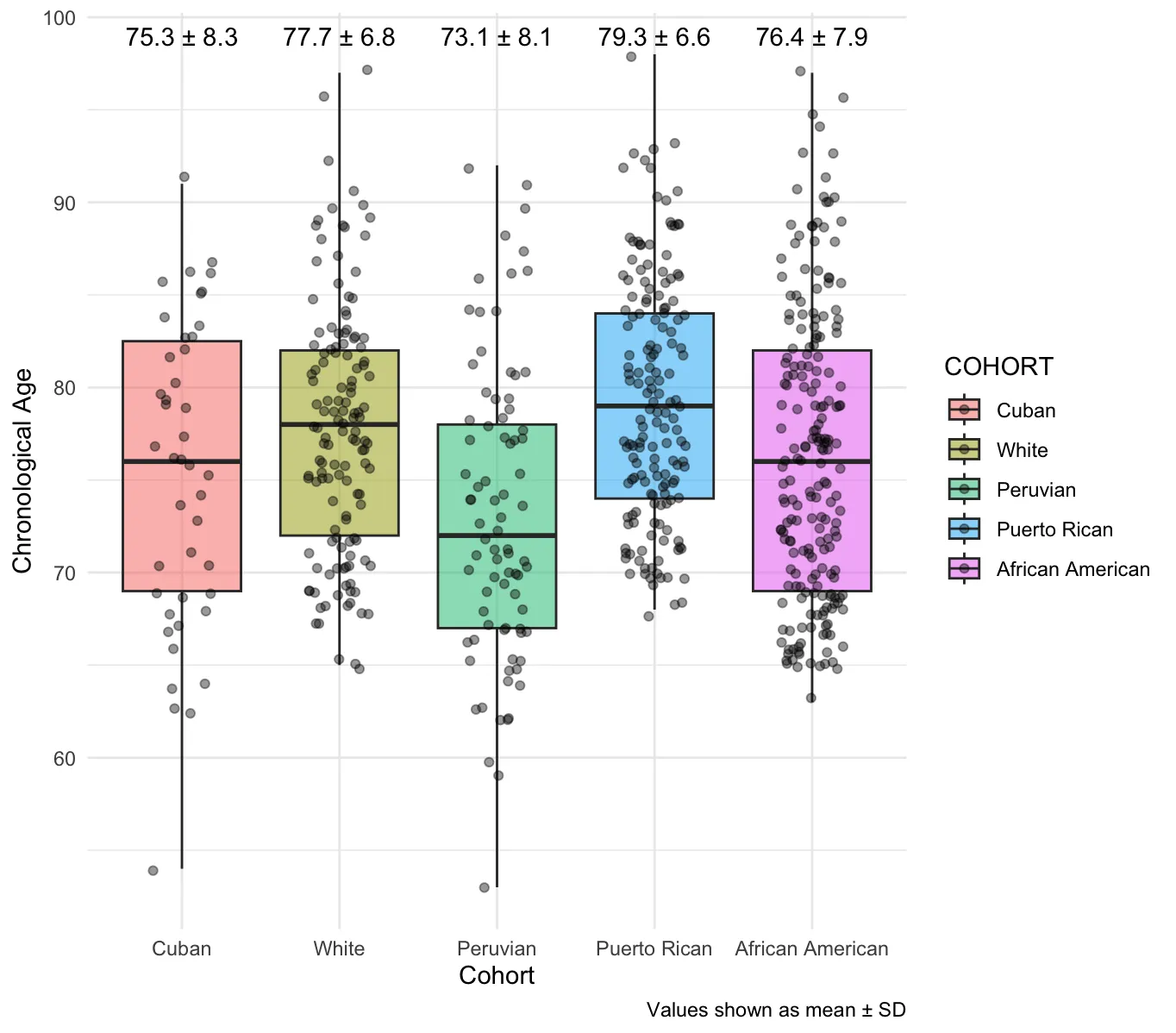

**Supplementary Figure 1: The age distributions of the MAGENTA study cohorts do not differ systematically.** We compiled the chronological ages for the individuals in the MAGENTA study and compared their distributions, stratifying by their respective cohorts. The differences in age distributions for the cohorts, particularly between Whites, Puerto Ricans, and African Americans, do not explain the significant difference in clock accuracy between these cohorts.

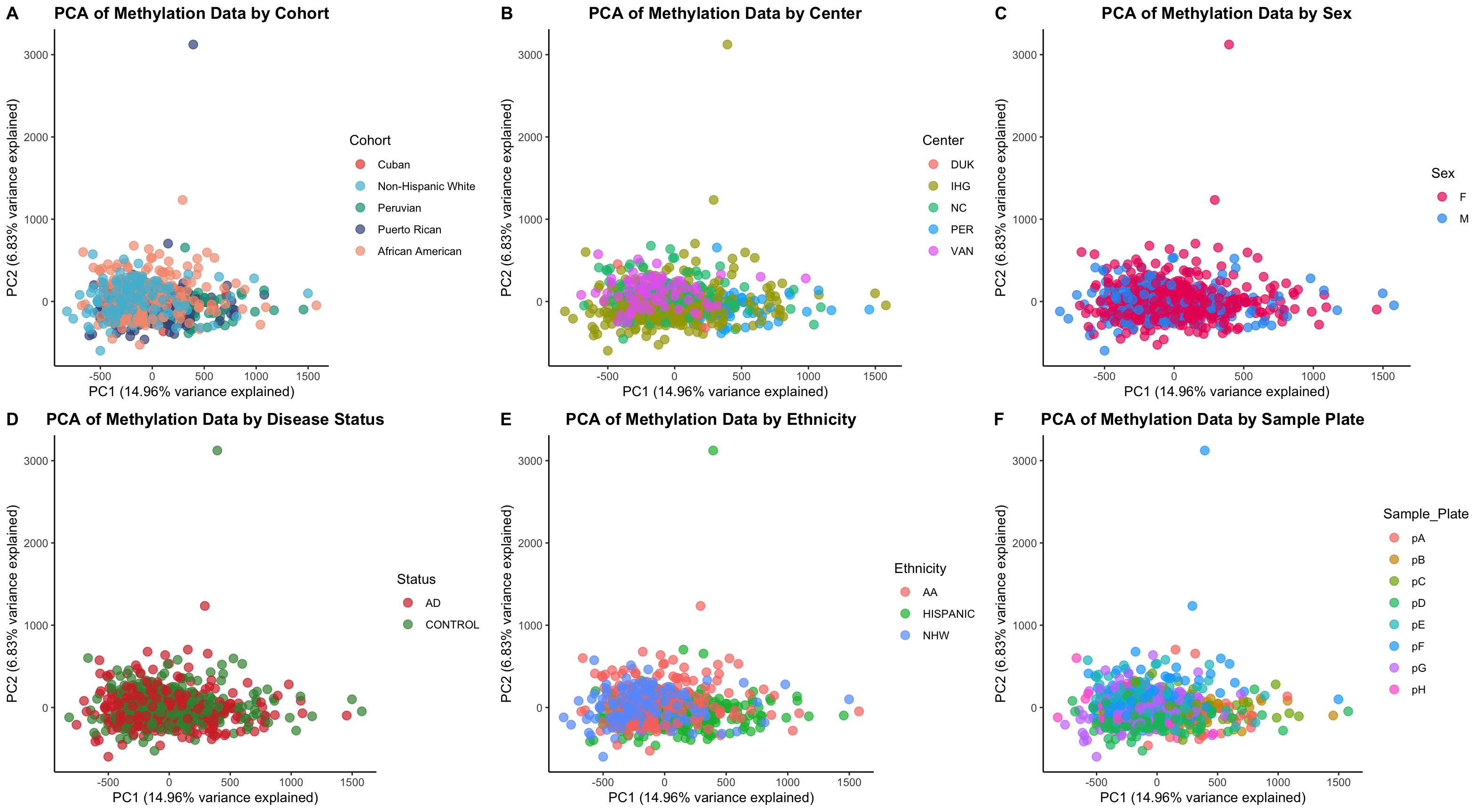

**Supplementary Figure 2: Principal component analysis of MAGENTA methylation data.** We performed a principal component analysis (PCA) of the normalized methylation data for all 621 MAGENTA individuals and calculated the variance explained by the first two PCs. PC1 explains 14.96% of the variance in the data, and PC2 explains 8.63% of the variance. Importantly, coloring the samples by cohort, sample center, sex, disease status, ethnicity, and sample plate did not show any outright stratification across the two PCs.

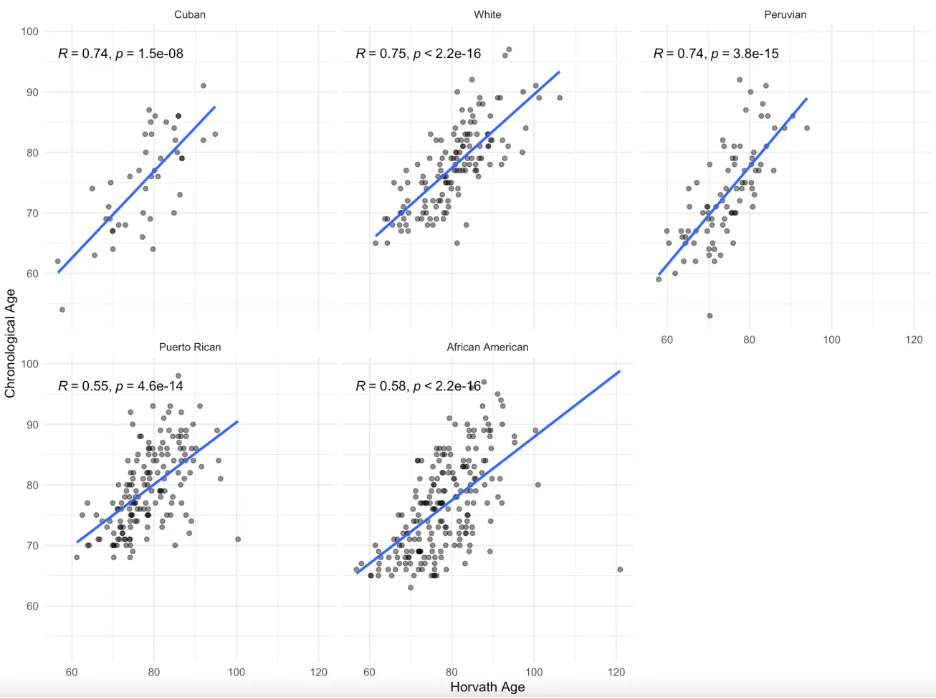

**Supplementary Figure 3: Combining cases and controls.** We calculated the correlation between Horvath DNAmAge and chronological age over combined AD cases and non-demented controls in the MAGENTA cohorts. The correlations did not change dramatically across the cohorts.

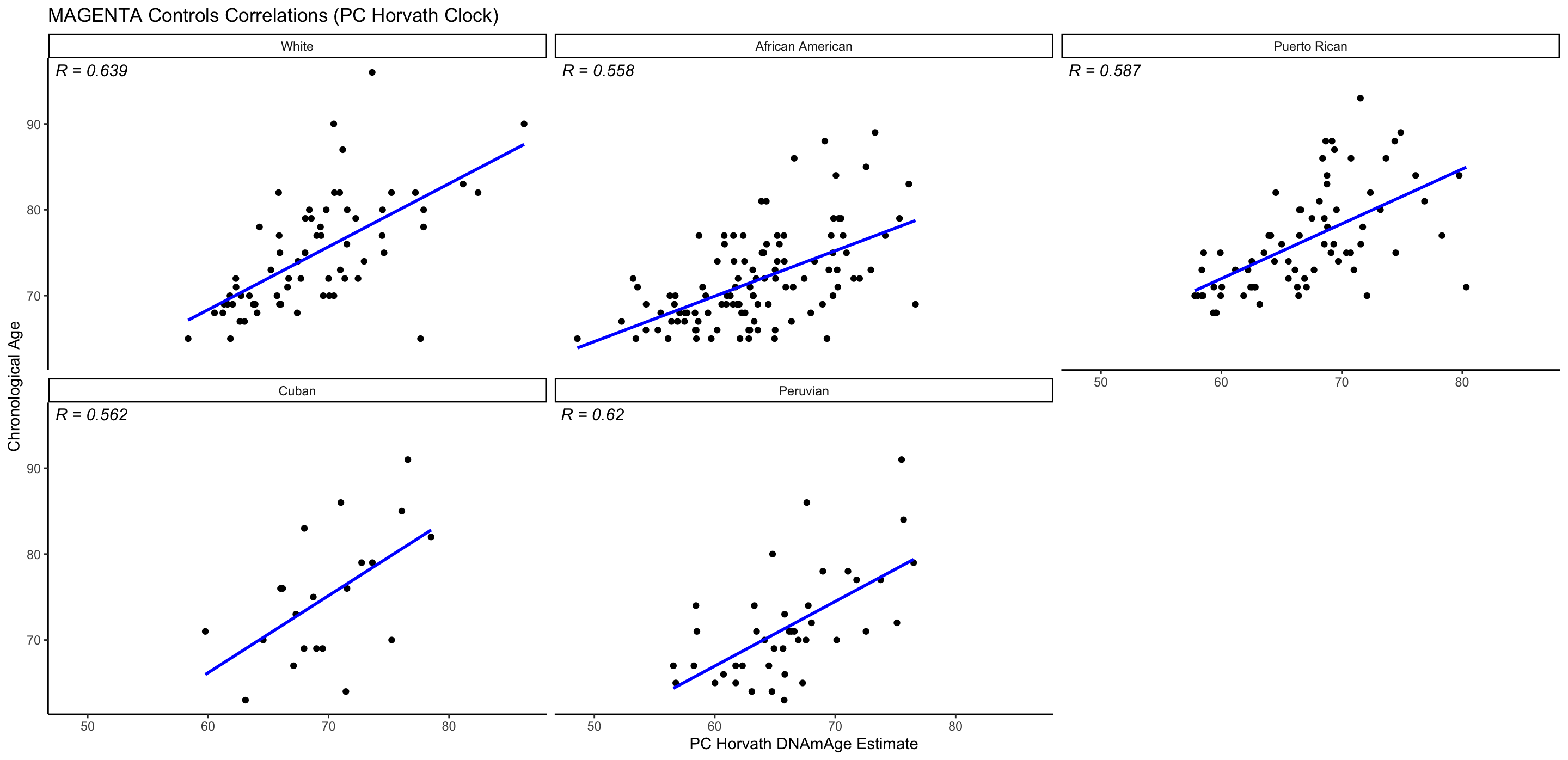

**Supplementary Figure 4: Principal component Horvath clock applied to MAGENTA controls.** We evaluated the correlation between the PC Horvath DNAmAge and chronological age in non-demented controls from the MAGENTA cohorts. This version of the clock did not yield consistent improvements in age prediction accuracy or generalizability across MAGENTA cohorts.

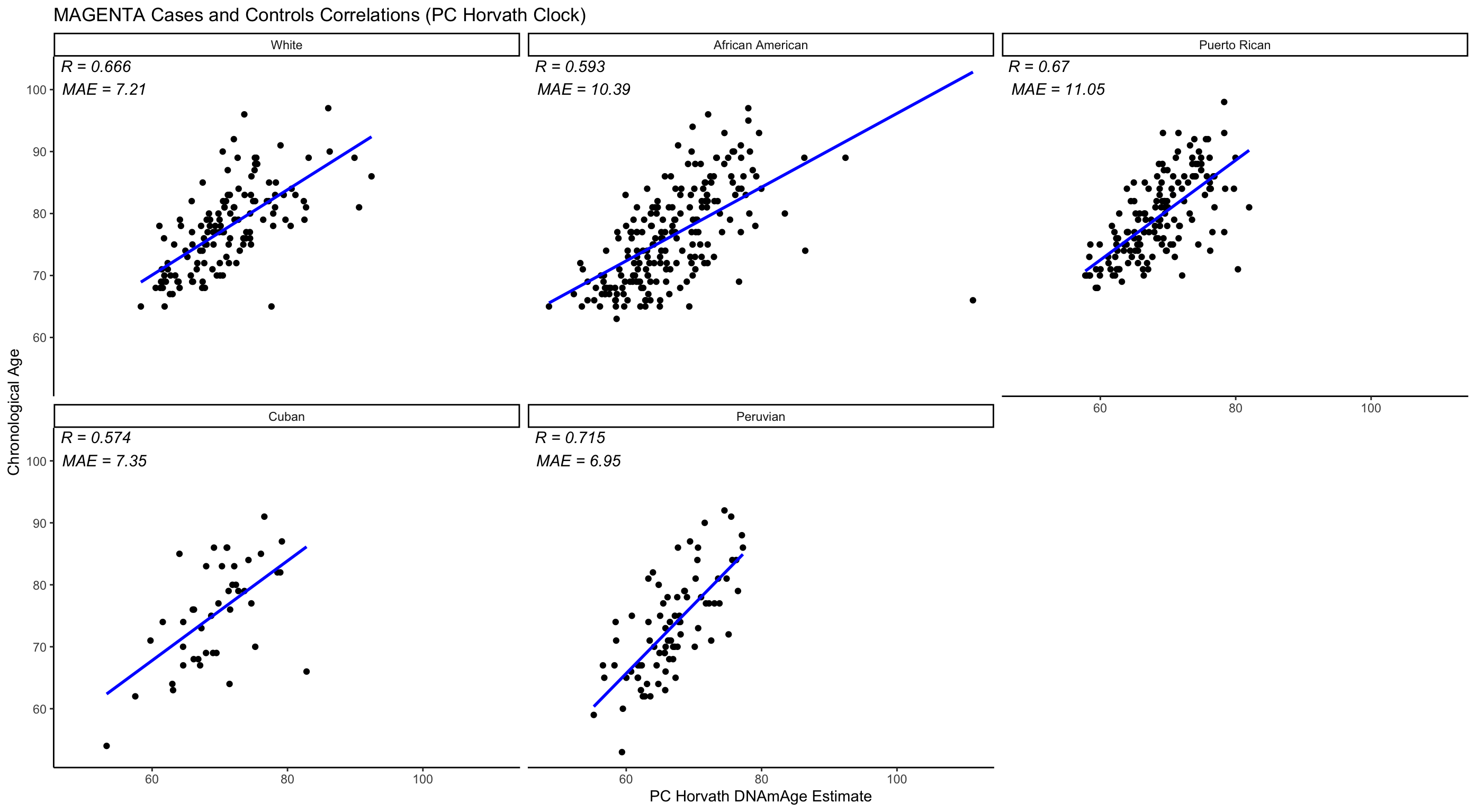

**Supplementary Figure 5: Principal component Horvath clock applied to MAGENTA cases and controls.** We evaluated the correlation between the PC Horvath DNAmAge and chronological age in both cases and non-demented controls from the combined MAGENTA cohorts. This approach did not consistently improve the clock’s performance compared to the controls alone.

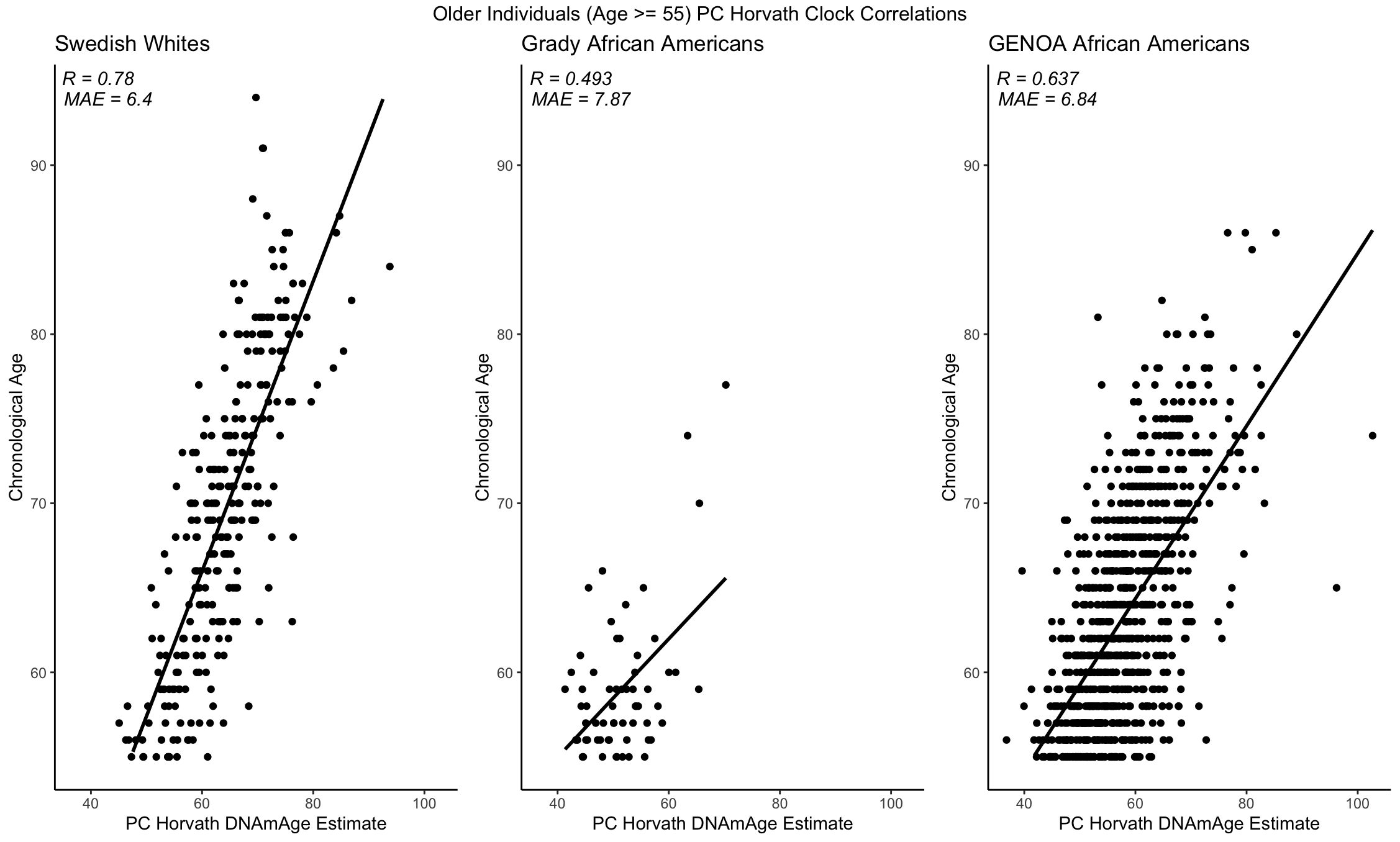

**Supplementary Figure 6: Principal component Horvath clock applied to replicate datasets.** We evaluated the correlation between the PC Horvath DNAmAge and chronological age in the three external, replication datasets of White Swedish individuals and African Americans. We observed results consistent with those obtained on the MAGENTA cohorts.

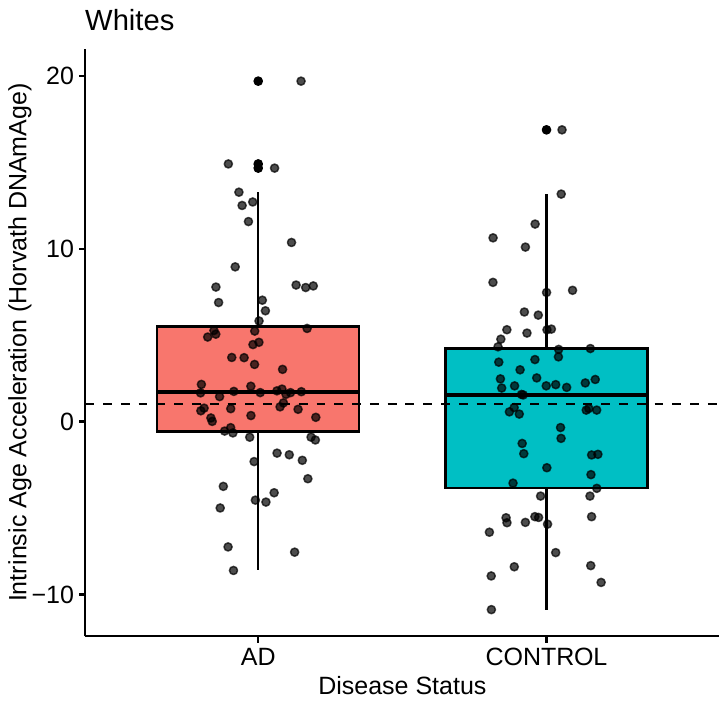

**Supplementary Figure 7: The Horvath Clock accurately discerns between white AD patients and non-demented controls.** Horvath clock intrinsic age acceleration was significantly greater on average for white AD patients than matched, non-demented controls (median 1.7 vs. 1.5 years, p = 0.041), consistent with previous literature.

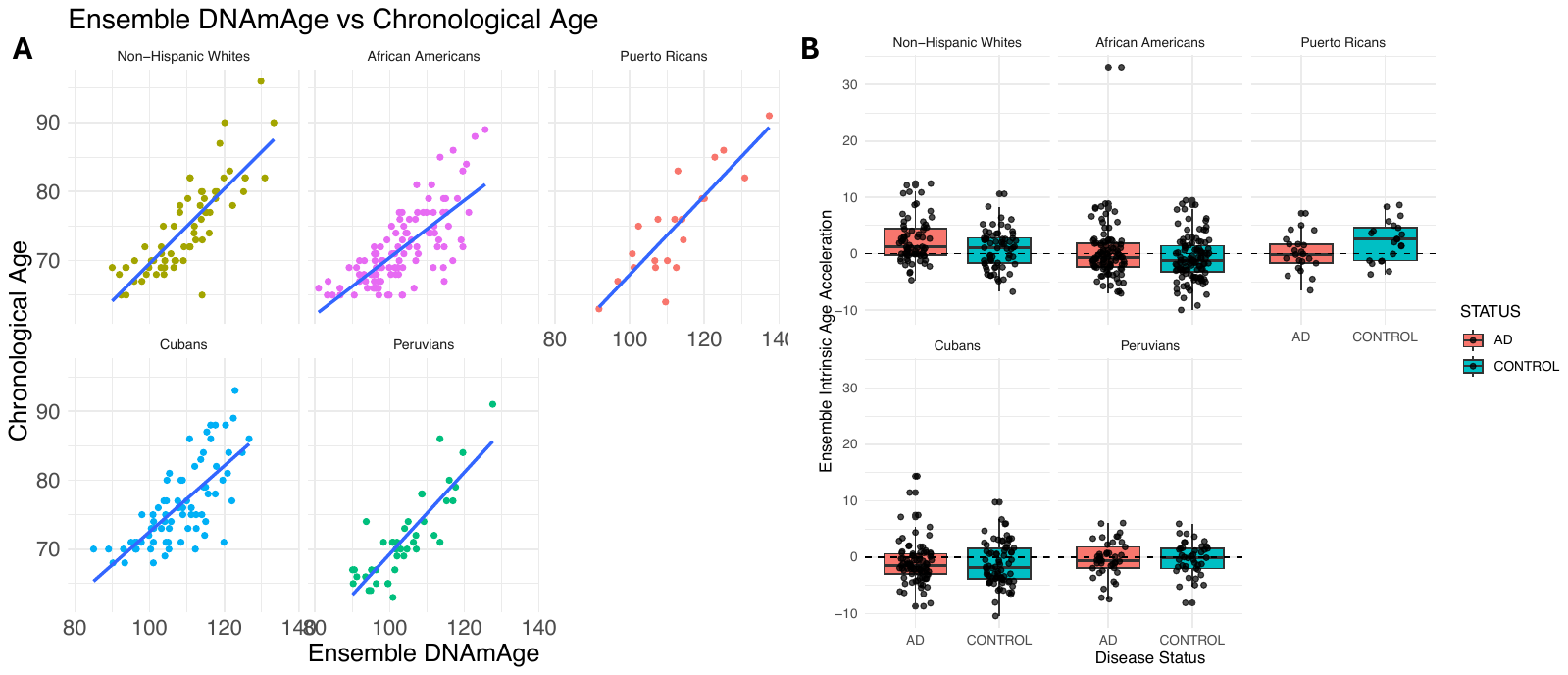

**Supplementary Figure 8: Combinations of DNAmAge and intrinsic age accelerations from multiple methylation clocks do not improve their performance.** We combined multiple methylation clocks and their respective predictions for all MAGENTA individuals but saw no increase in accuracy over individual clocks.

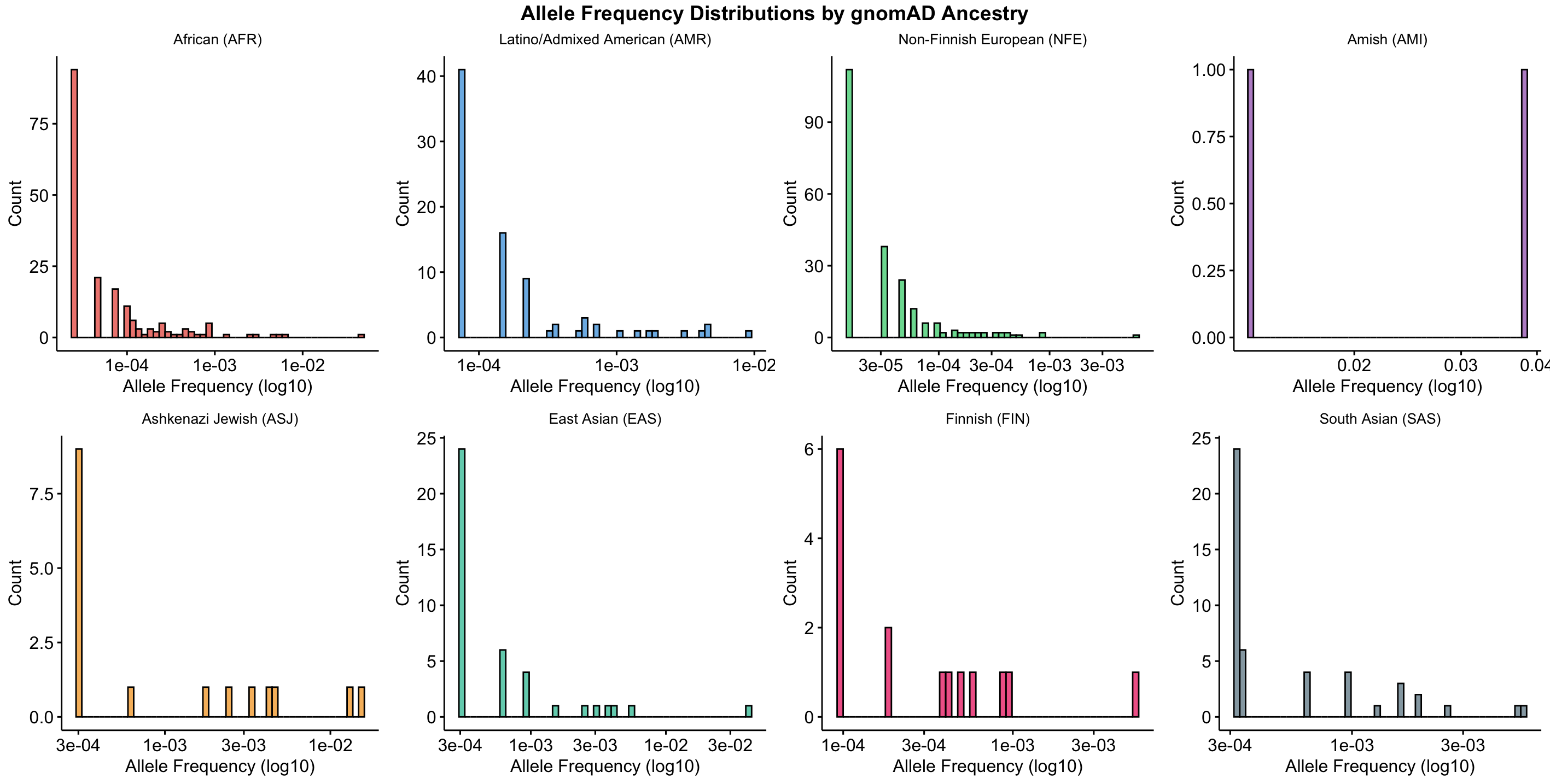

**Supplementary Figure 9: Allele frequencies for Horvath clock CpG-disrupting variants across each gnomAD v.3.0 ancestry.** The allele frequencies of the clock CpG-disrupting variants were very low, even after stratification by gnomAD ancestries. Note that the Amish had only two variants in the Horvath clock CpGs, but only one was common at 1% allele frequency.

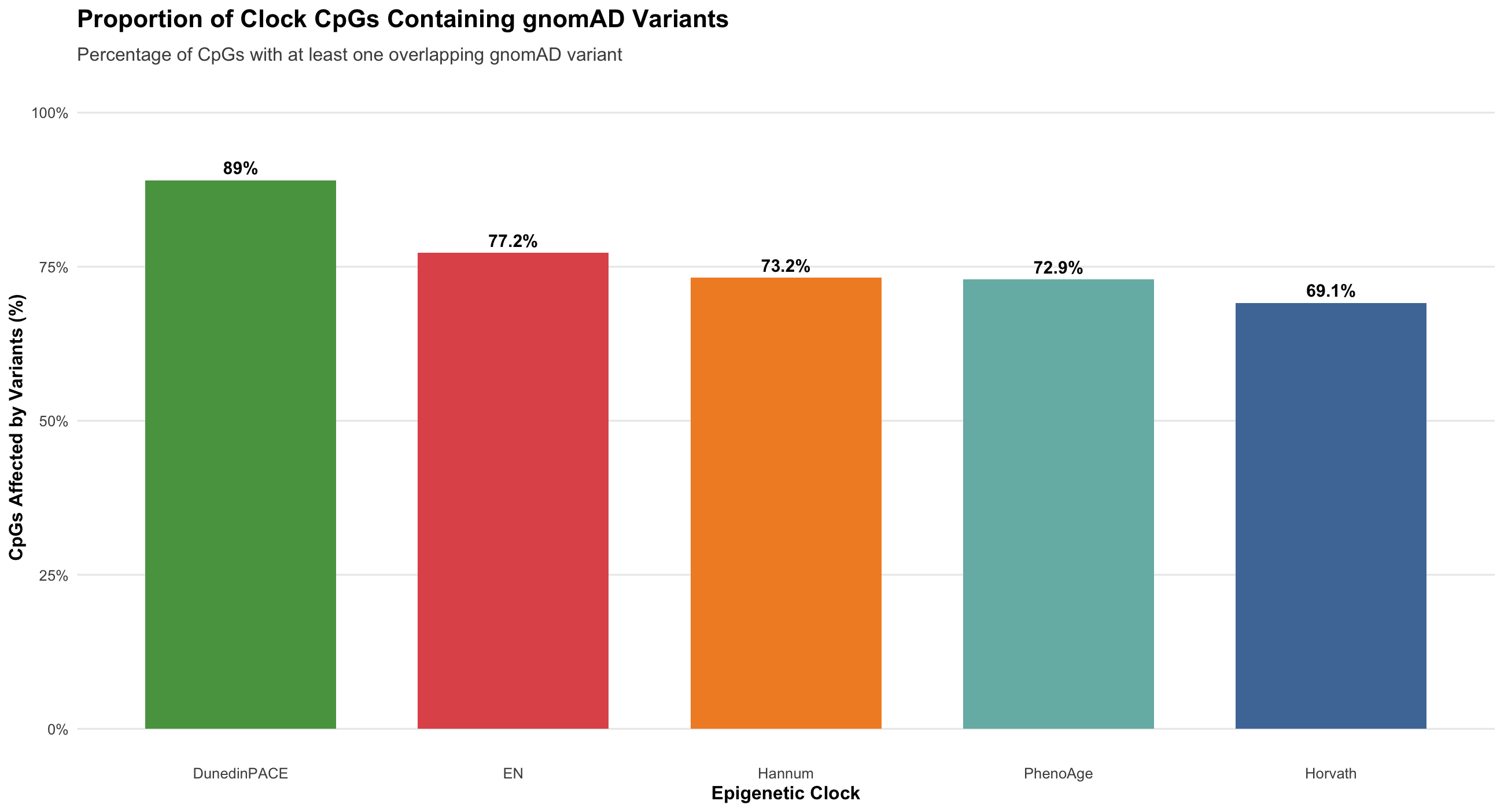

**Supplementary Figure 10: Overlap of gnomAD version 4.1 variants and clock CpG sites across multiple clocks.** We intersected the genomic coordinates of all clock CpG sites across all clocks tested in our study with all variants in the latest version of the gnomAD database. Many variants overlap the clock CpG sites, but only a small number for each clock are common (AF > 1%): Hannum (1), Zhang19_EN (5), PhenoAge (2), and DunedinPACE (1).

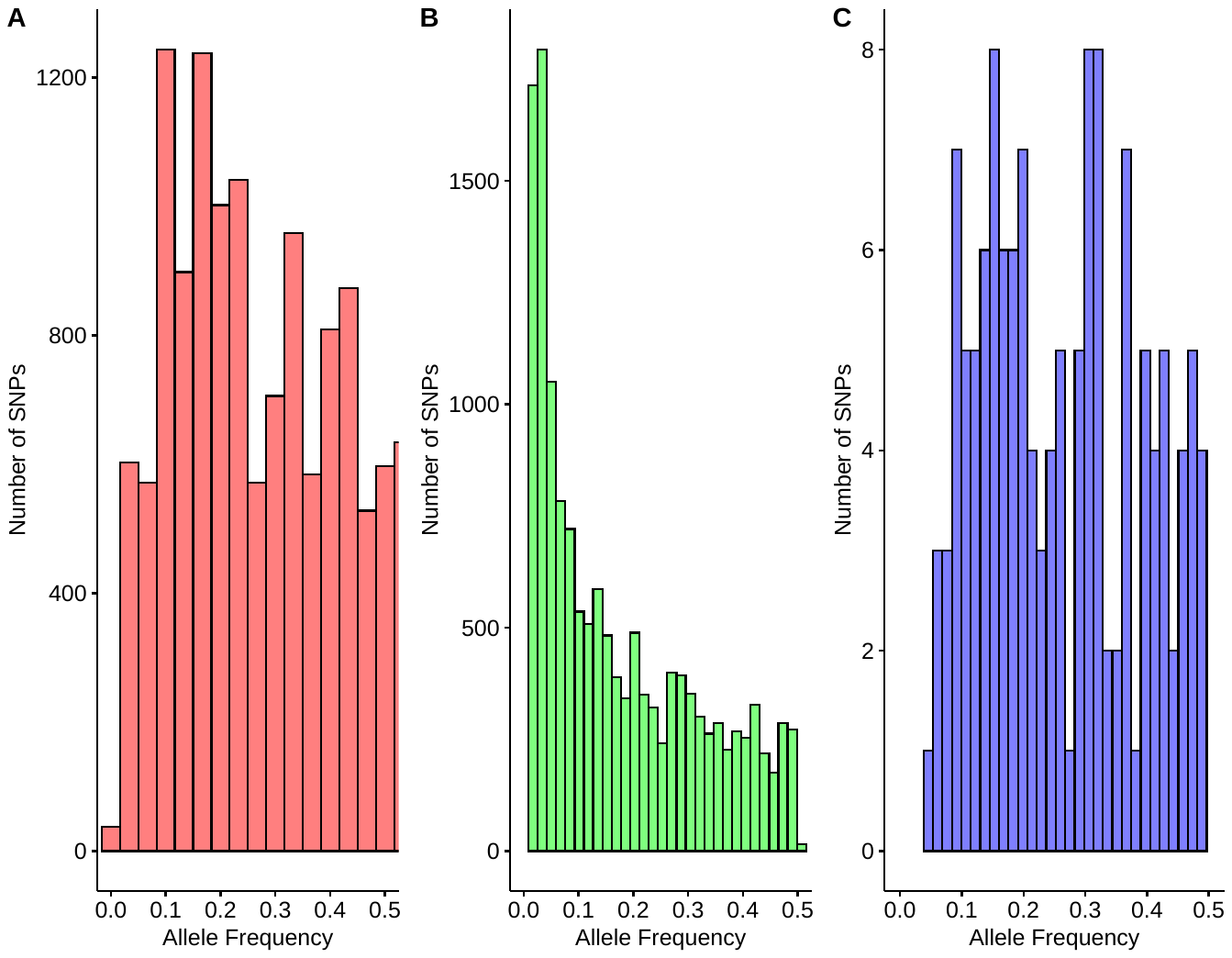

**Supplementary Figure 11: Allele frequencies of meQTL that affect Horvath clock CpG sites**. We compiled the reported allele frequencies for thousands of variants that affect the methylation levels of Horvath clock CpG sites. Most variants are reported to be common in their respective study populations. **Panel A**: variants from EUR and SAS individuals; **Panel B**: variants from African American individuals; **Panel C**: variants from EUR individuals in the UK.

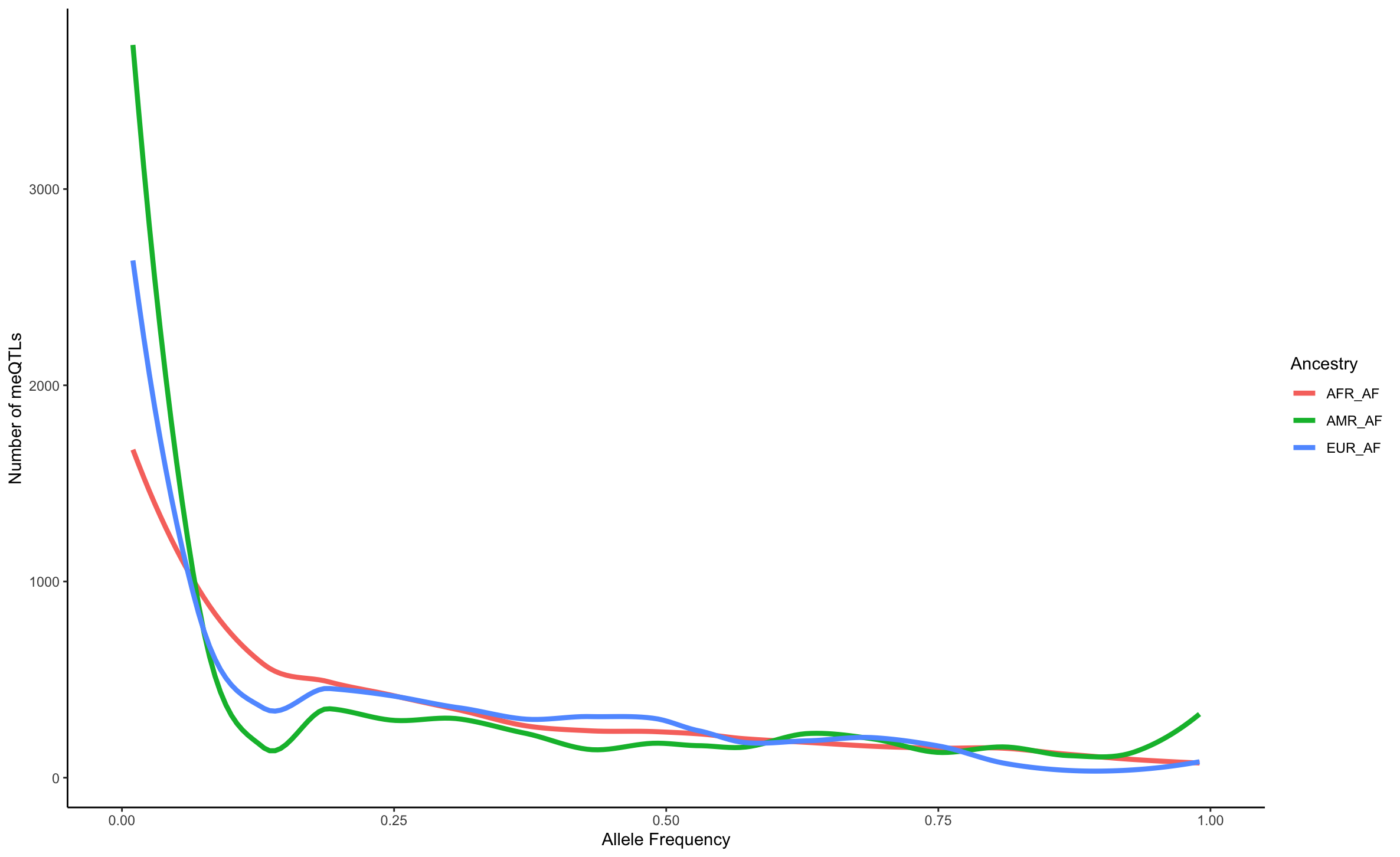

**Supplementary Figure 12: Allele frequencies of meQTL that affect Horvath clock CpG sites, stratified by local ancestry blocks in gnomAD admixed individuals**. The allele frequency distribution of the 17,180 unique variants associated with methylation levels at Horvath clock CpGs found in gnomAD Latino admixed individuals. Clock meQTL have significantly higher allele frequencies in African local ancestry genomic segments among the 7,612 Latino admixed individuals with varying proportions of European, American, and African ancestry from gnomAD v3.1.2.

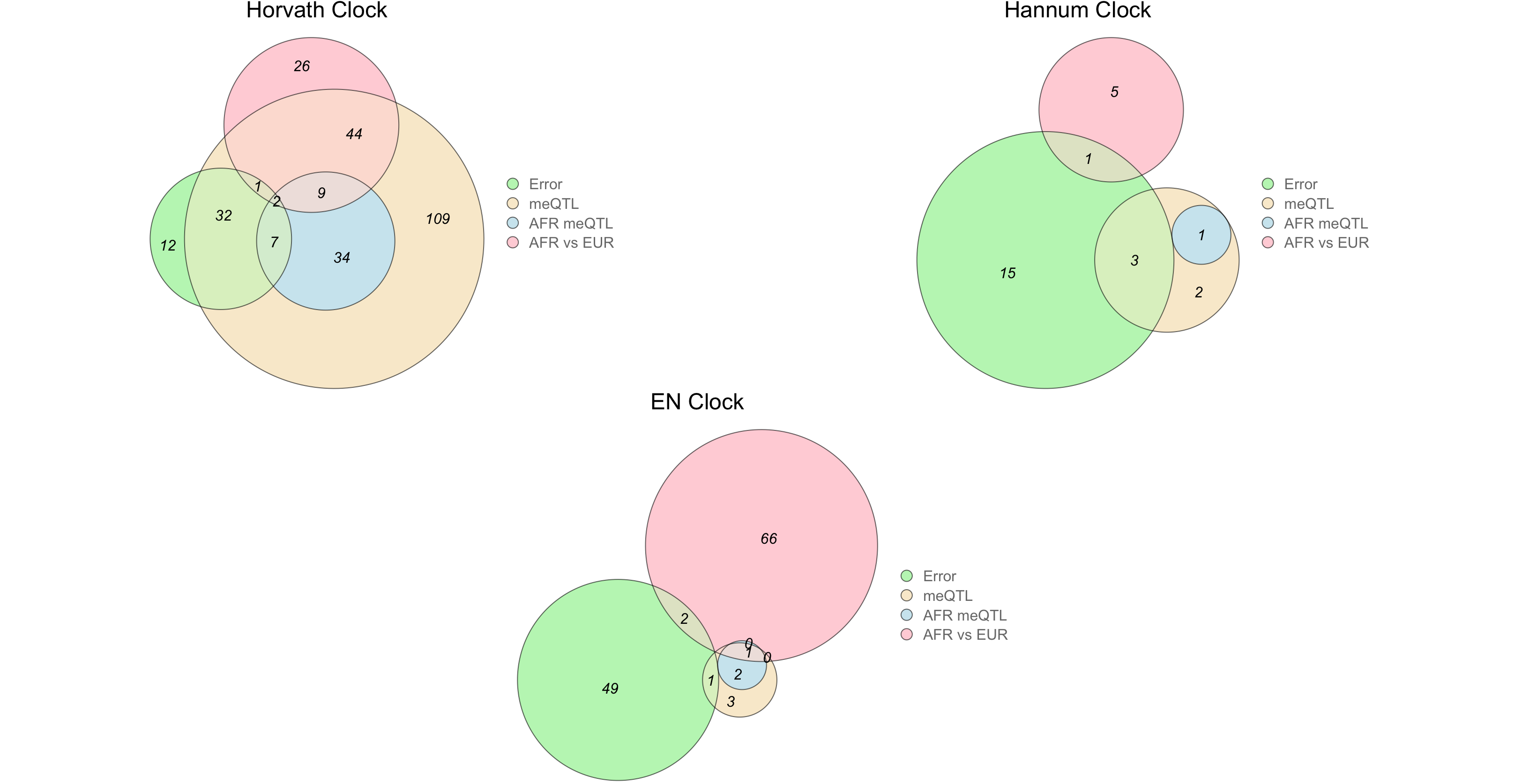

**Supplementary Figure 13: Venn diagrams for overlap of clock CpG sites.** The four-way overlap includes sets of clock CpGs whose methylation levels are significantly associated with increased clock error in MAGENTA individuals, clock CpGs that are differentially methylated in whole blood samples of African ancestry individuals relative to whole blood samples of European ancestry individuals (adjusting for chronological age and sex), clock CpGs affected by meQTL, and clock CpGs affected by African ancestry-differentiated meQTL.

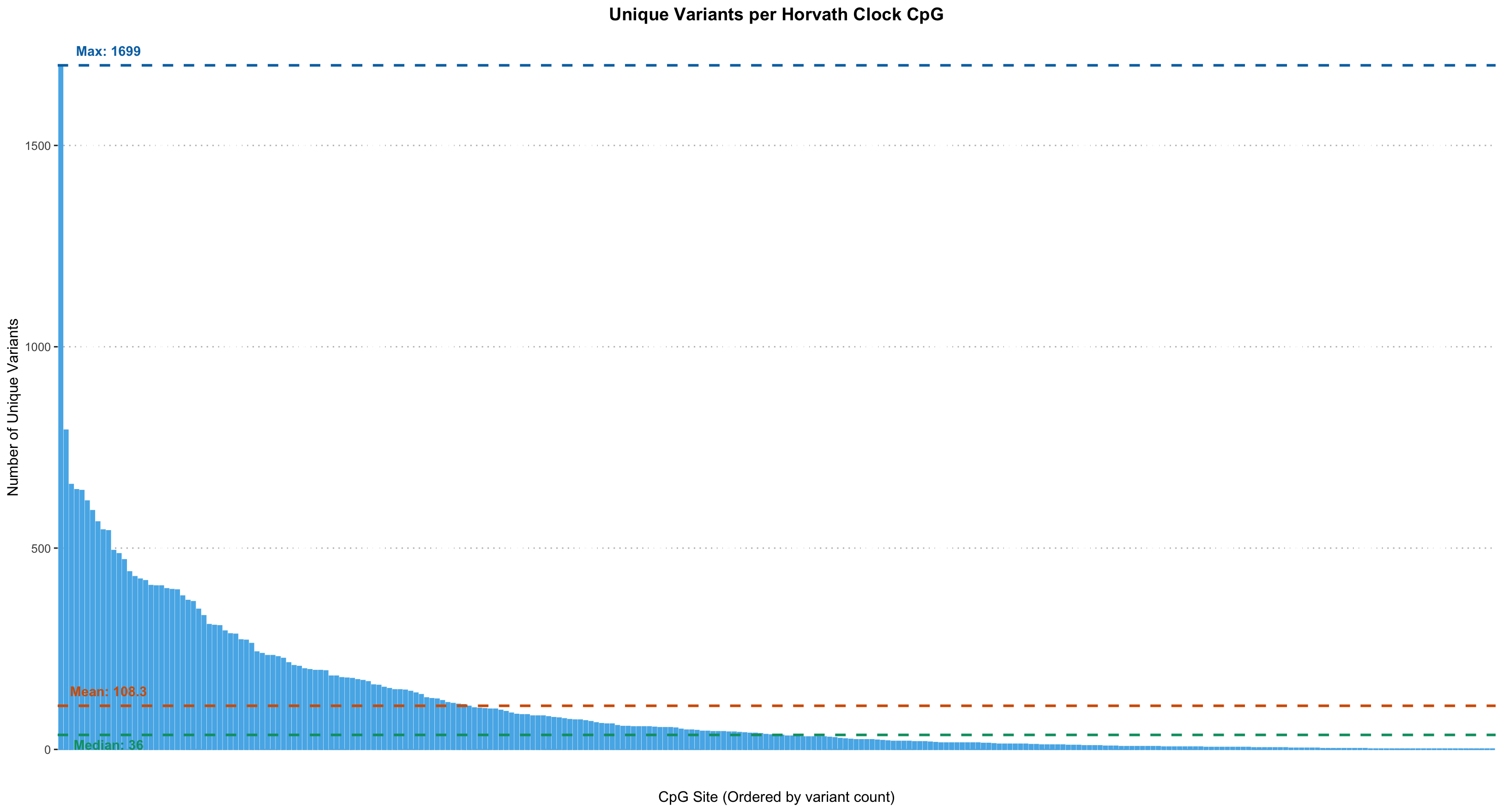

**Supplementary Figure 14: Distribution of unique meQTL affecting Horvath clock CpG sites.** We analyzed the distribution of the meQTL from EUR, SAS, and AFR individuals that affect Horvath clock CpG sites. Blue dashed lines indicate the maximum number of variants affecting a clock CpG site (1,699), orange dashed lines indicate the mean (108.3), and green dashed lines indicate the median (36).

**Supplementary Tables**

**Supplementary Table 1: Performance metrics for the Horvath Clock applied to non-demented controls in the MAGENTA cohorts.**

| MAGENTA Cohort | Sample Size | Median Absolute Error (MAE) | Mean Squared Error (MSE) | R |
| --- | --- | --- | --- | --- |
| Whites | 65 | 5.10 | 38.7 | 0.72 |
| African Americans | 107 | 5.38 | 46.7 | 0.51 |
| Puerto Ricans | 74 | 5.19 | 48.4 | 0.45 |
| Peruvians | 41 | 4.19 | 26.8 | 0.72 |
| Cubans | 21 | 5.60 | 53.6 | 0.68 |

**Supplementary Table 2: Summary of Horvath Clock performance across all evaluated cohorts (individuals** $\boldsymbol{\geq}$**55 years old).**

| Cohort | Sample Size | Median Absolute Error (MAE) | Mean Squared Error (MSE) | R |
| --- | --- | --- | --- | --- |
| MAGENTA Whites | 133 | 4.90 | 38.6 | 0.75 |
| MAGENTA African Americans | 205 | 5.62 | 60.4 | 0.58 |
| MAGENTA Puerto Ricans | 158 | 4.92 | 42.7 | 0.55 |
| MAGENTA Peruvians | 82 | 4.68 | 33.1 | 0.74 |
| MAGENTA Cubans | 43 | 5.18 | 42.7 | 0.74 |
| Grady Project African Americans | 62 | 5.49 | 53.4 | 0.57 |
| GENOA Study African Americans | 865 | 4.12 | 39.7 | 0.67 |
| White Swedish Individuals | 280 | 3.73 | 34.9 | 0.79 |
